# Gliotoxin, identified from a screen of fungal metabolites, disrupts 7SK snRNP, releases P-TEFb and reverses HIV-1 latency

**DOI:** 10.1101/848929

**Authors:** Mateusz Stoszko, Abdullah M.S. Al-Hatmi, Anton Skriba, Michael Roling, Enrico Ne, Yvonne M. Mueller, Mohammad Javad Najafzadeh, Raquel Crespo, Joyce Kang, Renata Ptackova, Pritha Biswas, Alessia Bertoldi, Tsung Wai Kan, Elisa de Crignis, Robert-Jan Palstra, Miroslav Sulc, Joyce H.G. Lebbink, Casper Rokx, Annelies Verbon, Wilfred van Ijcken, Peter D. Katsikis, Vladimir Havlicek, Sybren de Hoog, Tokameh Mahmoudi

## Abstract

A leading pharmacological strategy towards HIV cure requires “shock” or activation of HIV gene expression in latently infected cells with Latency Reversal Agents (LRAs) followed by their subsequent clearance. In a screen for novel LRAs we used fungal secondary metabolites (extrolites) as a source of bio-active molecules. Using orthogonal mass spectrometry (MS) coupled to latency reversal bioassays, we identified gliotoxin (GTX) as a novel LRA. GTX significantly induced HIV-1 gene expression in latent ex vivo infected primary cells and in CD4+ T cells from all aviremic HIV-1+ participants. RNA sequencing identified 7SK RNA, the scaffold of the P-TEFb inhibitory 7SK snRNP complex to be significantly reduced upon GTX treatment of independent donor CD4+T cells. GTX disrupted 7SK snRNP, releasing active P-TEFb, which then phosphorylated RNA Pol II CTD, inducing HIV transcription. Our data highlight the power of combining a medium throughput bioassay, mycology and orthogonal mass spectrometry to identify novel potentially therapeutic compounds.

## Introduction

Combination anti-retroviral therapy (cART) causes a drastic and immediate viral decrease by targeting distinct steps in the HIV-1 life cycle effectively blocking replication and halting disease progression (Ho et al., 1995; Perelson et al., 1997; Wei et al., 1995). However, cART does not target or eliminate HIV that persists in a latent state in cellular reservoirs. Because some of the proviruses are replication competent, latent HIV infected cells inevitably rebound once c-ART is interrupted, leading to necessity for life-long therapy (Siliciano and Siliciano, 2015). Particularly in resource-limited countries, which are also disproportionally affected, this is translated into an insurmountable medical, social and financial burden. To achieve a scalable cure for HIV infection, it will be necessary to reduce or eliminate the latent HIV infected reservoir of cells and/or equip the immune system with the robustness and effectiveness necessary to prevent viral rebound such that c-ART can be safely discontinued.

An important breakthrough in HIV-1 cure was the unequivocal proof that it is possible to mobilize the latent patient HIV reservoir by treatment with agents that activate HIV gene expression (LRAs) (Archin et al., 2012). However, clinical studies thus far have shown little to no reduction in the latent reservoir in patients (Spivak and Planelles 2017; Rasmussen and Søgaard 2018). This is consistent with limited potency and specificity of currently tested drugs, which appear to be unable to reach a significant proportion of latently infected cells, or to induce HIV-1 expression in latent reservoir at sufficient levels to produce viral proteins for recognition by the immune system (Kim et al., 2018). Furthermore, transcriptional stochasticity and heterogeneity of latent HIV integrations (Battivelli et al., 2018) may pose additional barriers to reactivation of the latent reservoir as a whole; sequential rounds of stimulation yield new infectious particles (Ho et al., 2013), while certain LRA combinations result in more efficient latency reversal when administered in intervals rather than at once (Bouchat et al., 2016). In addition, pleiotropic functions and toxic effects of LRAs may compromise the ability of CD8+ T cells to eliminate HIV protein expressing cells (Jones et al., 2014; Zhao et al., 2019). Therefore, it is critical to identify and develop novel therapeutics, which strongly induce HIV-1 gene expression to effectively disrupt HIV latency without dampening the immune response.

The pharmaceutical industry is highly equipped for high throughput screens using defined synthetic libraries. While this is an effective approach, it is important to record that approximately half of the novel small molecules introduced to the market between 1981 and 2014 are natural or nature-derived (Newman and Cragg, 2016). Biological systems represent an invaluable source of functional molecules with high chemical diversity and biochemical specificity, evolved during millions of years of adaptation (Cary et al., 2016; Richard et al., 2018; Vo and Kim, 2010; Wang et al., 2017; Yasuhara-Bell et al., 2010). In particular, fungi represent a largely unexplored source of compounds with potential therapeutic use. Fungi secrete a gamut of extracellular compounds and other small-molecular extrolites (Sanchez et al., 2012). While some of these compounds have been shown to have antibiotic (ex. penicillin) or carcinogenic (ex. aflatoxin) properties, little is known in general about their biological activities and possible molecular targets. In addition, a single fungal strain often produces a wide array of secondary metabolites which are not essential for its growth but are exuded as a consequence of specific environment such as nutrient-rich *versus* minimal growth conditions (Brakhage, 2013; Přichystal et al., 2016). Fungal extrolites might target various signaling pathways in mammalian cells, such as those influencing HIV-1 gene expression. Fungal supernatants are an ideal source for an expert academic setting where low and medium throughput biological screening systems, academic knowledge of evolutionary mycology, and state-of-the-art fractionation and purification techniques are routinely combined.

Studies of regulation of HIV-1 gene expression have identified distinct molecular mechanisms and cellular pathways at play, which can be targeted pharmacologically to activate expression of latent HIV (De Crignis and Mahmoudi, 2016; Ne et al., 2018; Stoszko et al., 2019). The rich diversity of fungal extrolites therefore, may prove an untapped source of new compounds that target HIV for reactivation. In search of novel LRAs, here we have performed an unbiased medium through-put screen of fungal extrolites, coupled to medium through-put HIV latency reversal bioassays and orthagonal fractionation and mass spectrometry-NMR and identified Gliotoxin (GTX). GTX potently reversed HIV-1 latency in multiple in vitro latency models as well as ex vivo in cells obtained from all aviremic HIV-1 infected patients examined without associated cytotoxicity. We demonstrate that GTX disrupts the P-TEFb-sequestering 7SK snRNP complex, leading to degradation of its scaffold 7SK RNA, and release of active P-TEFb, which is then recruited by Tat to phosphorylate the Ser2 residue of the RNA Pol II C-Terminal Domain (CTD) and activate HIV transcription elongation.

## Results

### Growth supernatant of *Aspergillus fumigatus* identified in a medium through-put screen of fungal secondary metabolite possesses HIV-1 latency reversal activity

We screened 115 species of filamentous fungi for their ability to induce HIV-1 proviral expression; of species that appeared promising, 2-4 additional strains were tested (Supplementary Table 1). Species belonged to 28 orders (43 families) of the fungal Kingdom (Fig. 1a) and were chosen based on their evolutionary position, ecological trends, and known active production of extracellular compounds. The majority of fungi were of ascomycetous affinity, 4 species were of basidiomycetous affinity and two belonged to the lower fungi. Selected fungi were grown in both complete yeast media as well as minimal media (RPMI), as they are known to produce distinct extrolites depending on their growth conditions (Supplementary Fig. 1). Culture supernatants were then screened for latency reversal activity using Jurkat derived 11.1 and A2 cell line models of HIV-1 latency (J-Lat) in a low-medium through-put assay set up, in which expression of GFP is controlled by the HIV-1 promoter and indicates latency reversal. We identified the supernatant of *Aspergillus fumigatus* CBS 542.75 to strongly activate the latent HIV-1 5’LTR (Fig. 1b). We also compared other *Aspergillus* species growth supernatants indicated for their potential to induce HIV-1 expression (Fig. 1c) and observed that only strains of *A. fumigatus* (CBS 542.75, CBS 113.26 and CBS 100074) possessed latency reversal activity (Fig. 1c).

**Fig. 1.**
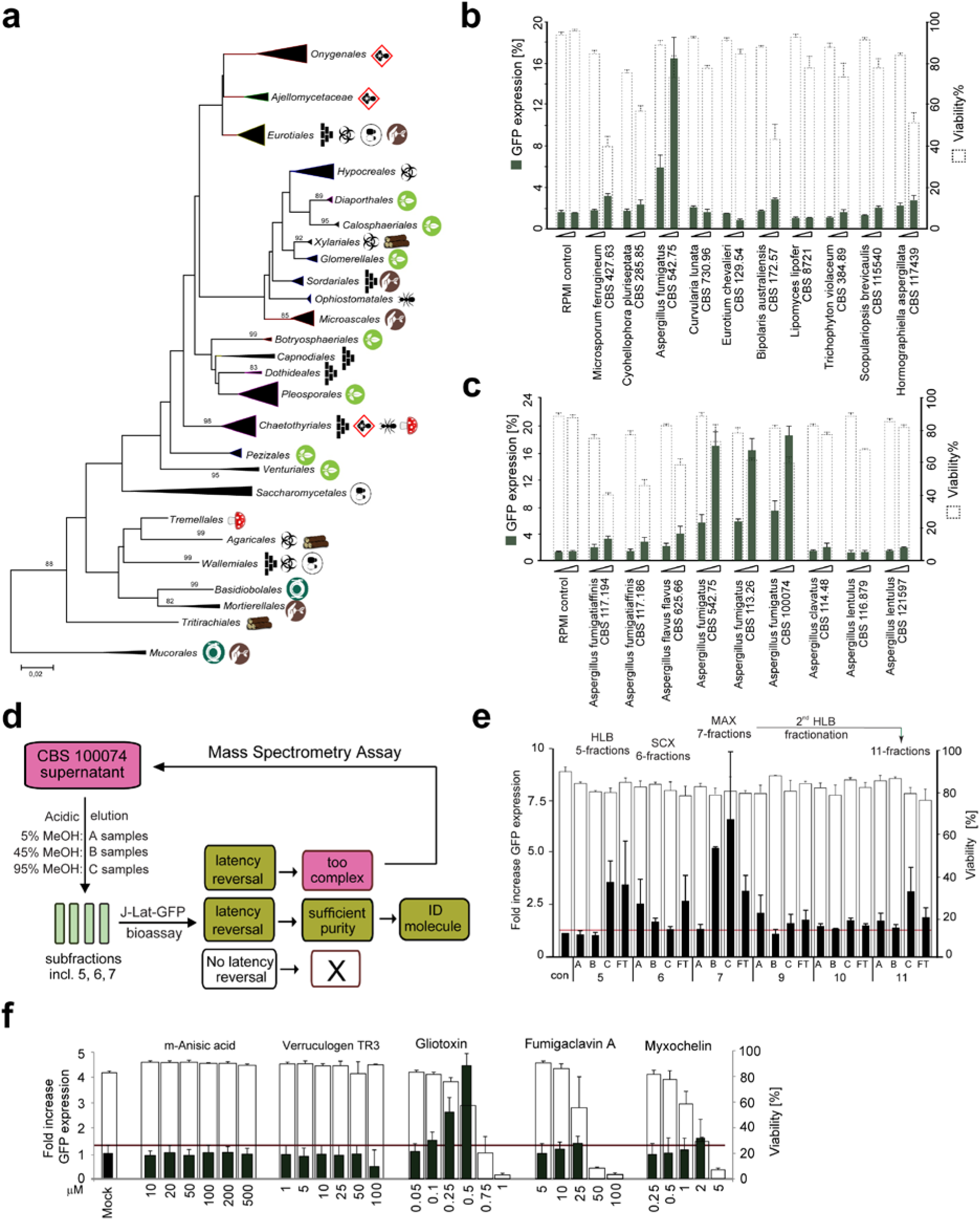
Medium through-put screen of fungal secondary metabolites combined with orthogonal fractionation and MS strategy coupled to latency reversal bioassays identifies Gliotoxin from growth supernatant of *Aspergillus fumigatus* to reverse HIV-1 latency. **a,** Phylogenetic tree representing main orders of the fungal Kingdom with strains used in current study, collapsed per order. Orders selected from the tree published by Gostincar et al. (2018), with some of the lower orders included for structural reasons. Approximate ecological trends in the orders are summarized by symbols, as follows: 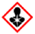 vertebrate pathogenicity prevalent, 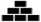 climatic extremotolerance prevalent; 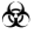 frequent production of extracellular metabolites or mycotoxins, 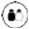 frequent osmotolerance or growth in sugary fluids, 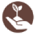 numerous members with soilborne lifestyle, 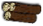 numerous members inhabiting decaying wood rich in hydrocarbons, 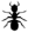 frequent insect-association, 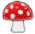 frequent mushroom decomposition or hyperparasitism on fungi or lichens, 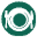 frequent inhabitants of foodstuffs or vertebrate intestinal tracts. **b,** Latency reversal bioassay performed by treatment of J-Lat A2 cells with increasing volumes of growth (normalized by O.D.) supernatants obtained from selected fungal strains. **c,** Latency reversal bioassay in J-Lat A2 cells with growth supernatants obtained from members of the *Aspergillus* genus. Cells were treated as in panel B. **d**, Schematic representation of the orthogonal MS strategy coupled to latency reversal bioassays used to identify putative LRA. See main text for full description. **e,** Three pre-concentration cartridges (HLC, SCX, MAX) were combined with variable content of extracting solvent (A: 5% MeOH, B: 45% MeOH, C: 95% MeOH, FT = Flowthrough). Latency reversal potential of fractionated secondary fungal metabolites was tested via treatment of J-Lat A2 cells. Latency reversal (%GFP, left axis, black bars) and cell viability (%viable, right axis, empty bars) was assessed by flow cytometry analysis. **f,** Commercially obtained versions of five common molecules identified in active fractions were tested for LRA activity in J-Lat A2 cells. Data are presented as fold increase in GFP expression as indicated, ± SD from at least three independent experiments.

### Orthogonal LC-MS/NMR strategy coupled to latency reversal bioassays identifies Gliotoxin (GTX) from growth supernatant of *Aspergillus fumigatus* as a putative LRA

Due to the chemical complexity of the positive fungal supernatants, direct mass spectrometry (MS) analysis of their constituents proved to be impossible. Therefore, *A. fumigatus* CBS 100074 growth supernatant was fractionated several times by means of orthogonal MS (Fig. 1d). We selected this particular supernatant as it showed the highest potency to reverse latency in the J-Lat models. After each round of fractionation, all samples/fractions were again tested in latency reversal bioassays, followed by quantitation of the GFP expression and identification of fractions retaining latency reversal activity. As expected, originally less active fractions became more active during the fractionation/enrichment process (Fig. 1e). The most active 7B/7C fractions were further fractioned on HLB cartridge (11-samples) and components of 7B/7C and 11C fractions de-replicated by Cyclobranch software (Fig. 1e and Supplementary Fig. 2a) (Novák et al., 2017). Compound matching against the annotated database of *Aspergillus* secondary metabolites revealed a set of candidate compounds further selected for latency reversal testing (Supplementary Fig. 2b and Supplementary Table 2). Among candidate molecules identified, Gliotoxin (GTX), obtained from a commercially available synthetic source, was able to induce expression of the latent pro-virus in a concentration dependent manner (Fig. 1f). Of note, GTX was found in all positive fractions (Supplementary Table 2). Interestingly, while GTX isolated from supernatant of CBS 100074 showed strong induction of HIV-1 transcription, supernatant of *Aspergillus flavus* (CBS 625.66), a close relative of *A. fumigatus* which was inactive in latency reversal (Fig 1c), did not contain GTX (Supplementary Fig. 2c and d), providing further support that GTX is the main mediator of LRA activity in *A. fumigatus* supernatants.

### GTX reverses latency in ex vivo infected primary CD4+ T cells without associated cytotoxicity

To examine the latency reversal potential of GTX in a more clinically relevant system, we employed a modified primary *ex vivo* infected latency model in which primary CD4+ T cells are infected with a full length replication incompetent HIV-1 virus driving expression of the luciferase reporter (Supplementary Fig. 3a) (Lassen et al., 2012). Treatment of latently infected CD4+ T cells with GTX resulted in significant, concentration-dependent latency reversal at lower concentrations than necessary to achieve re-activation in latently infected cell-lines (Fig. 2a-c). GTX [20 nM] treatment resulted in over 20-fold induction of HIV-1 expression (Fig. 2a), which translated into over 10% latency reversal of that observed upon maximal stimulation with PMA/Ionomycin (Fig. 2b). Similar to the latency reversal observed using commercially available synthetic GTX (Fig. 2a-b), treatment of HIV-1 infected latent primary CD4+ T cells with GTX [20 nM] isolated from growth supernatant of Aspergillus fumigatus CBS 100074 resulted in significant latency reversal, approximately 10% of maximal PMA/ionomycin stimulation (Fig. 2c). At high concentrations GTX is known to be toxic to immune cells (Stanzani et al., 2005; Suen et al., 2001; Yamada et al., 2000), ascribed to its unusual disulfide bridge, responsible for pleiotropic effects on cellular and viral systems (Scharf et al., 2016). Consistent with the literature, GTX concentrations upward of 100 nM induced significant toxicity as indicated by Annexin V staining (Fig 2d and Supplementary Fig. 3b). However, primary CD4+ T cell viability was not significantly affected after GTX treatment at concentrations up to 50 nM (Fig 2d and Supplementary Fig 3b). Thus, at lower concentrations in which GTX did not show toxicity on CD4+ T cells, strong latency reversal was induced (Figs. 2a-d and Supplementary Fig. 3b). CD8+ T cells play a central role in eliminating HIV-infected cells (Trautmann, 2016). Therefore, it is of utmost importance to evaluate potential toxicity of newly developed LRAs on CD8+ T cells (Zhou et al., 2019). Importantly, GTX at a low concentration of 20 nM did not reduce the viability of CD8+ T cells whether unstimulated or αCD3/αCD28-stimulated PBMCS were examined (Figs. 2e-f, Supplementary Fig. 4a-b). Consistent with the literature (Hur et al., 2008; Orciuolo et al., 2007; Stanzani et al., 2005; Suen et al., 2001; Sutton et al., 1995; Wichmann et al., 2002; Yamada et al., 2000; Zhou et al., 2000), treatment with higher concentrations of GTX at 100 nM and 1 µM caused apoptosis and death of primary CD4+ and CD8+ T cells as well as B cells, NK cells and monocytes (Figs. 2d-e, 3c and Supplementary Figs. 3b, 4a-c). Potential for clinical applicability of a candidate LRA also requires that it does not induce global T-cell activation, nor should it interfere with CD8+ T cell activation. Treatment of unstimulated primary CD4+ T-cells and CD8+ T cells with GTX [20 nM], which significantly reversed latency, did not induce expression of the T cell activation markers CD69 and CD25 (Fig. 2g, 3d, Supplementary Fig. 5a-b), nor did it induce proliferation of resting CD4+ and CD8+ T cells (Supplementary Fig. 5c-d), while, as expected, PMA/Ionomycin treatment activated T cells (Fig. 2g). Conversely, GTX treatment of activated PBMCs also did not inhibit CD25 expression (Fig. 3d, Supplementary Fig. 5b) or proliferation of activated CD4+ or CD8+ T cells (Supplementary Fig. 5c-d).

**Fig. 2.**
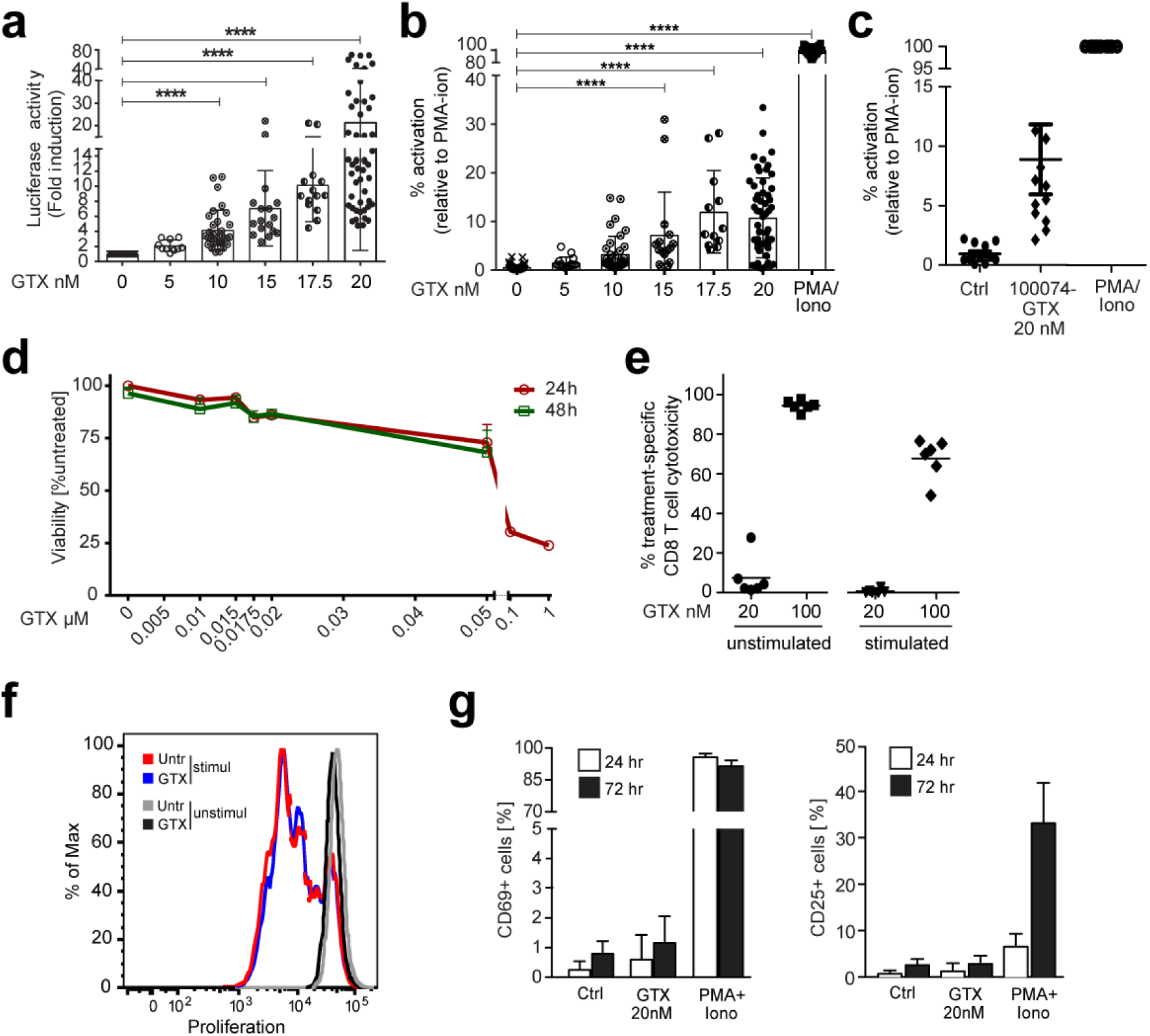
GTX [20 nM] reverses HIV-1 latency in ex vivo infected primary CD4+ T cells without associated cytotoxicity, T cell activation, or inhibition of proliferative capacity. **a**, Latency reversal after treatment of latently infected primary CD4+ T cells with increasing concentrations of GTX for 24 hours as indicated shown as fold induction over untreated control luciferase activity. **b**, Data presented in **a** shown as percent latency reversal activity normalized and shown as percent activity relative to treatment with the positive control PMA/Ionomycin. **c,** Latency reversal after treatment of HIV-1 infected latent primary CD4+ T cells with [20nM] GTX isolated from growth supernatant of *Aspergillus fumigatus* CBS 100074 measured as increased luciferase activity normalized to and shown as percent of activation of the positive control PMA/Ionomycin. Experiments were performed in duplicate using cells obtained from at least 6 healthy donors. Wide horizontal lines represent average, shorter horizontal lines represent standard deviation. **d,** Viability of the primary CD4+ T cells treated for 24 (red line) and 48 (green line) hours with indicated increasing concentrations of GTX, assessed by FACS on the basis of forward versus side scatter. **e,** Unstimulated or αCD3/αCD28 stimulated PBMCs were treated with GTX as indicated for 72 h followed by Annexin V staining of CD8+CD3+ T cells and flow cytometry. Each symbol represents one healthy donor (n = 6 from 3 independent experiments using 2 different donors cells), horizontal line depicts mean. **f,** Representative FACS plot overlay showing the division of unstimulated and αCD3/CD28-stimulated CD8+ T cells in the presence or absence of GTX. **g,** Activation status of primary CD4+ T cells upon GTX treatment as indicated. Averaged data of 3 independent experiments performed using 2 different donors cells in duplicate (n=6).

**Fig. 3.**
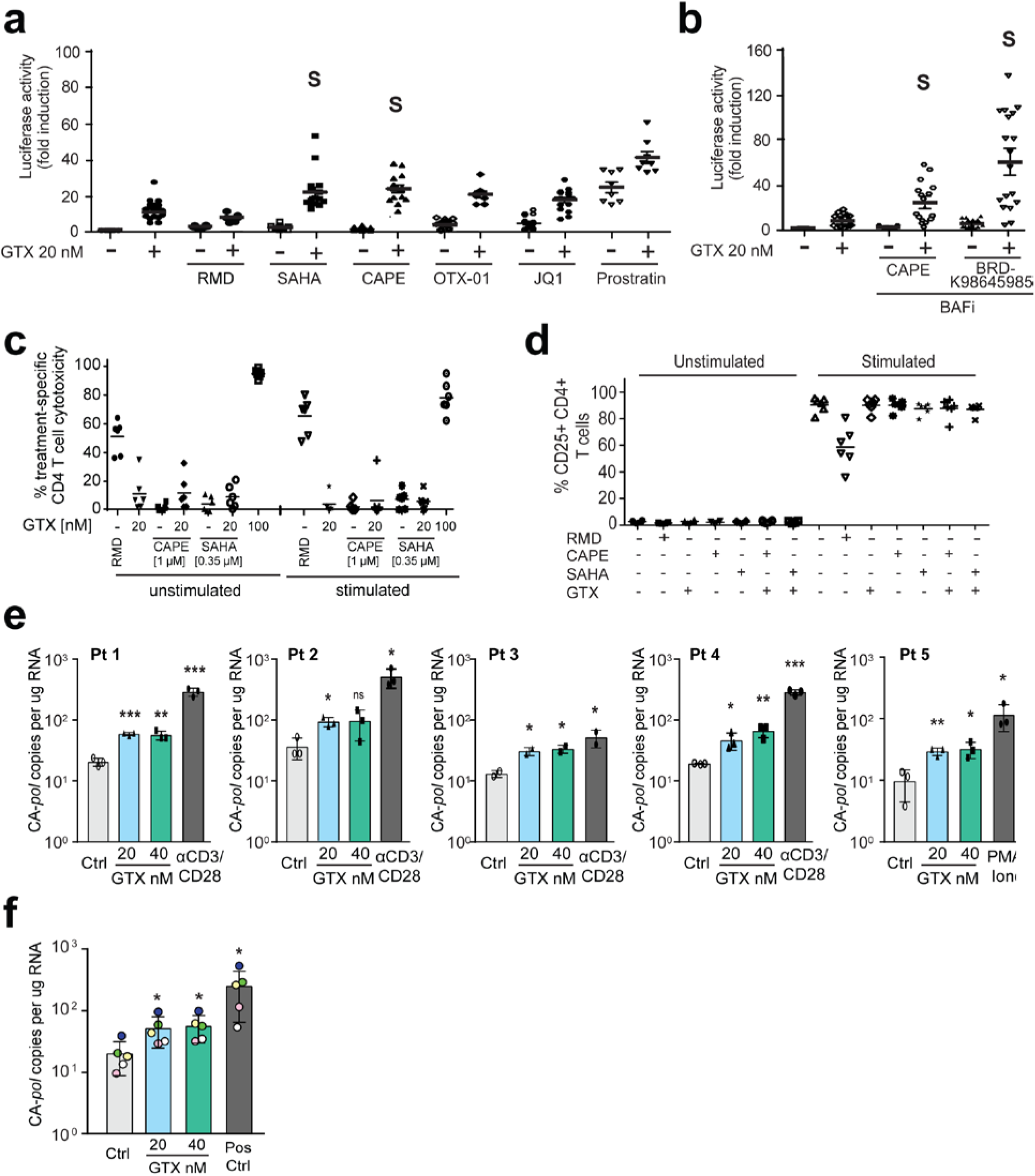
GTX strongly synergizes with HDAC and BAF inhibitors and reverses latency in primary CD4+ T cells obtained from all tested aviremic participants. **a-b**, HIV-1 latency reversal in latent ex vivo HIV-1 infected primary CD4+ T cells in response to 24 hour co-treatment with 20 nM GTX and distinct LRA class compounds as indicated and shown as fold increase in luciferase activity. S indicates compound synergism in latency reversal according to the Bliss independence score. **c,** Cytotoxicity of GTX alone and combined with indicated LRAs in CD4+ T cells. Unstimulated and αCD3/αCD28 stimulated PBMCs were co-treated as indicated for 72 h followed by Annexin V staining of CD4+CD3+ T cells. **d,** 20 nM GTX does not alter activation of CD4+ T cells. PBMC from healthy donors were incubated with the indicated LRAs for 72 hours either unstimulated or stimulated with αCD3/αCD28 antibodies. Panels depicts pooled data showing the frequency of CD25+ cells within CD4+ T cells. **e,** Absolute, cell associated (CA) *pol* copy number in CD4+ T cells isolated from 5 aviremic participants that were treated in vitro with vehicle control (Untr), GTX (20nM and 40nM) or positive control for 24 hours as indicated. Statistical significance was calculated using t test, * – p<0,05; ** – p<0,005; *** – p<0,0005. **f,** Data presented in panel A has been averaged and plotted together. Each symbol represents aviremic participant: green-participant 1; blue- participant 2; white- participant 3; yellow- participant 4; pink- participant 5. Statistical significance was calculated using unpaired Mann-Whitney test, * – p<0,05.

**Fig. 4.**
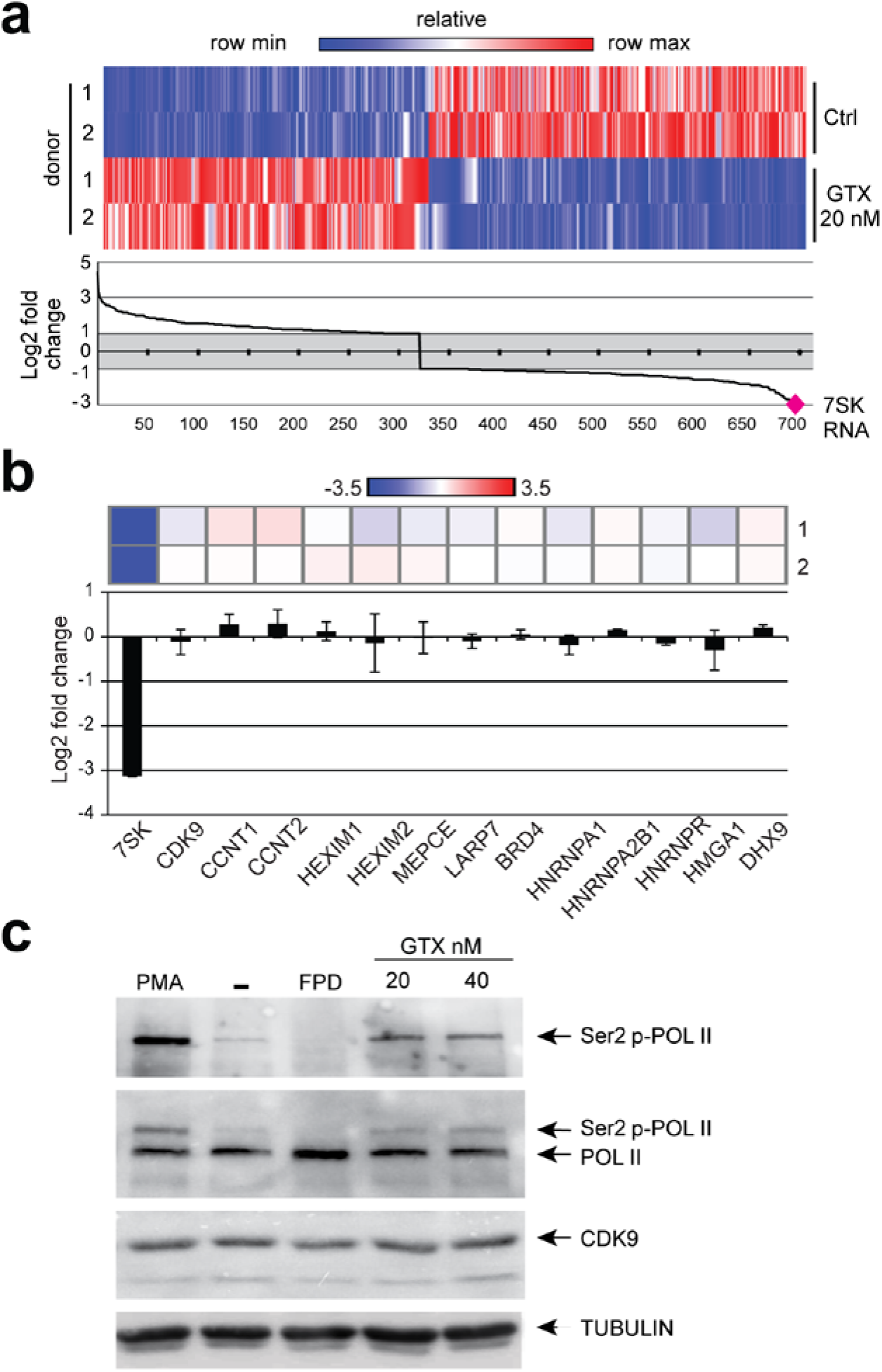
GTX treatment of resting CD4+ T cells causes decrease in 7SK RNA and activation of P- TEFb activity. **a,** Heat map of differentially expressed genes obtained from RNA sequencing analysis of primary CD4+ T cells treated as indicated for 4 h (top panel). RNA sequencing indicate 7SK RNA to be the most differentially decreased gene in response to GTX treatment of CD4+ T cells in two independent donors (bottom panel). **b,** GTX treatment of primary CD4+ T cells for 4 h leads to specific depletion of 7SK RNA levels and not mRNA levels of other components of the 7SK snRNP complex. **c,** Representative western blot analysis of CTD Serine 2 phosphorylation of RNA Polymarase II mediated by CDK9 in primary CD4+ T cells treated for 6h with GTX as indicated. PMA was used as a positive control, flavopiridol (FPD) was used as a negative control.

**Fig. 5.**
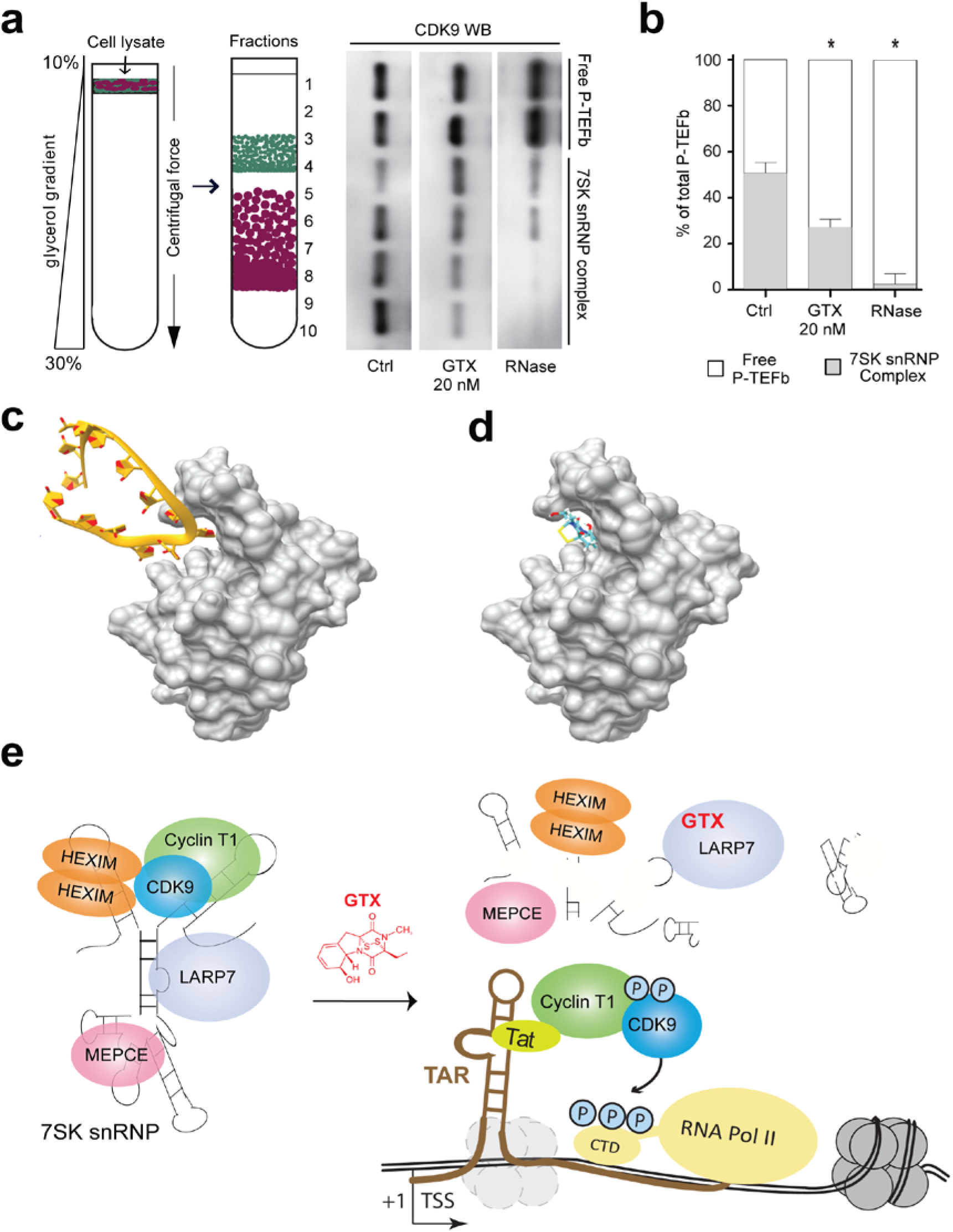
GTX disrupts 7SK snRNP, causing release of P-TEFb and enhanced HIV-1 transcription. **a,** GTX treatment causes release of pTEFb from sequestration within the 7SK snRNP complex. Left panel, schematic of glycerol gradient experiments in which cell lysates (from GTX treated or untreated cells) are loaded on top of generated glycerol gradients (10-30%) and ultracentrifuged, followed by collection of fractions, TCA protein precipitation and subsequent analysis by SDS-PAGE western blotting. Right panel, representative western blot analysis of glycerol gradient sedimentation of lysates from primary CD4+ T cell, which were treated (6h) with GTX [20 nM] or vehicle control as indicated using antibodies specific for the P-TEFb component CDK9. As control, untreated lysates were treated with RNAse for 1 hr to digest 7SK RNA and release P-TEFb. **b,** Quantification of free versus total CDK9 in primary CD4+T cells treated as indicated, as shown in **a** from three independent donors. Statistical significance was calculated using ANOVA followed by Tukey test (p < 0.01). **c-d,** Molecular docking of GTX onto LARP7. **c,** Crystal structure of human LARP7 C-terminal domain in complex with 7SK RNA SL4 (PDB ID code 6D12). LARP7 is shown in space-filling representation (grey) with bound RNA as yellow cartoon representation with bases indicated as flat rings (oxygens in red). One of the bases of 7SK RNA SL4 is bound into a deep pocket on the surface of LARP7. **d,** Proposed binding mode of GTX into the deep pocket on the surface of LARP7 as predicted by Chimera’s AutoDock Vina function and the Achilles Blind Docking Server. Color code for GTX is turquois for carbon, red for oxygen, yellow for sulphur, white for hydrogen, blue for nitrogen. **e,** Proposed model for GTX-mediated transcription activation of the latent HIV-1 LTR. GTX treatment leads to degradation of 7SK RNA, resulting in release of CDK9 (P-TEFb component) from the 7SK snRNP complex. Free P-TEFb is then recruited to the HIV-1 Tat-TAR axis, leading to phosphorylation of RNA polymerase II at Serine 2 and subsequent stimulation of transcription elongation.

### GTX enhances activity of other LRA class molecules and synergizes strongly with HDAC and BAF inhibitors

To investigate possible synergies, we tested the latency reversal potential of GTX [20 nM] in combination with a panel of known LRAs in the J-Lat A2 and 11.1 models of latency (Supplementary Fig. 6) as well as in *ex vivo* infected primary CD4+ T cells (Fig. 3a-b). GTX co-treatment enhanced the latency reversal activity observed after single treatments with all compounds (Fig. 3a-b and Supplementary Fig. 5d). Interestingly, when latent HIV-1 infected primary CD4+ T cells were co-treated with GTX [20 nM] and either the HDAC inhibitor SAHA or BAF inhibitors CAPE (Stoszko et al., 2015) or BRD-K98645985 (Marian et al., 2018), synergistic reversal of HIV-1 latency was observed (Fig. 3a-b). Co-treatments with BET inhibitors JQ-1 and OTX-015, as well as Prostratin resulted in an additive effect on HIV-1 pro-virus expression. Interestingly, in the primary CD4+ T cell model of HIV-1 latency, GTX treatment alone showed more potent latency reversal activity than SAHA, CAPE, OTX-015, JQ-1 or Romidepsin (RMD) alone at tested non-cytotoxic concentrations (Fig. 3a-b). RMD treatment showed modest latency reversal activity (Fig. 3a) and consistent with the literature (Zhao et al., 2019) significant CD4+ and CD8+ T cell cytotoxicity (Fig. 3c and Supplementary Fig 4a-b). With the exception of RMD, we did not observe any negative impact of these co-treatments on viability and proliferation of CD4+ T cells (Fig. 3c and Supplementary Fig. 5c), CD8+ T cells (Supplementary Fig. 4a-b, 5d) and other immune sub-populations including CD19+ B cells, CD56+ NK cells and CD14+ monocytes (Supplementary Fig. 4c). Moreover, none of the co-treatments altered the activation status of either resting or activated CD4+ and CD8+ T cells (Fig. 3d and Supplementary Fig. 5a-b). Our observed synergistic effects upon GTX co-treatment with BAF inhibitors and the HDAC inhibitor SAHA suggested that GTX may target a different pathway for HIV-1 latency reversal as targeting the same pathway with different molecules would more likely result in additive effects. Indeed, unlike SAHA, GTX treatment of CD4+ T cells did not result in increased histone acetylation (Supplementary Fig. 7a). Similarly, we excluded that GTX behaves as a PKC agonist, as treatment with GTX did not induce T cell activation, or the expression of PKC pathway target genes (Fig. 2g, Supplementary Fig. 7b).

### GTX reverses HIV-1 latency after ex vivo treatment of CD4+ T cells from all aviremic HIV-1 infected study participants

We next examined the potential of GTX to reverse latency after *ex vivo* treatment of CD4+ T cells obtained from people living with HIV-1. All five participants enrolled were treated with c-ART and maintained HIV-1 viremia below 50 copies/ml for at least two years. Despite differences in the size of the latent pool, assessed by maximal stimulation of the cells with α-CD3/CD28 beads or PMA/Ionomycin, GTX treatment at non-cytotoxic 20 and 40 nM concentrations significantly increased the levels of cell-associated HIV-1 *pol* RNA (CA-*pol*) in CD4+ T cells obtained from aviremic HIV-1+ participants (Fig. 3e-f). Notably, the observed GTX effect is systematic as latency reversal was uniformly observed in cells of all tested participants after 24 hours of stimulation (Fig. 3e-f). Additionally, we observed no increase in expression of genes related to T cell specific responses, reactive oxygen species or apoptosis, previously reported pleiotropic effects of GTX observed at significantly higher concentrations (Supplementary Fig. 7c). These results therefore indicate that GTX treatment at non-toxic concentrations reverses latency without inducing immune cell toxicity or activation and without affecting T cell proliferation, rendering GTX a promising potential candidate for further clinical investigation in context of HIV-1 latency reversal and inclusion in “shock and kill” strategies.

### GTX reverses HIV-1 latency via disruption of 7SK snRNP, leading to release of active P-TEFb

To gain insight into the molecular mechanism by which GTX reverses latency, we performed RNA sequencing of primary resting CD4+ T cells isolated from healthy blood donors that were treated for 4 hours with 20 nM GTX. This short incubation time was chosen to focus on primary effects of GTX on the global transcriptome, decreasing the presence and likelihood of secondary transcriptional effects. We observed a very good correlation between treatments of two independent healthy blood donors (Fig. 4a top panel). Moreover, less than 700 genes showed an altered differential expression pattern (Fig. 4a bottom panel). Interestingly, small nuclear RNA 7SK was the most decreased (more than 9-fold) transcript after GTX treatment in CD4+ T cells from the independent donors (Fig 4a). 7SK RNA serves as a scaffold for the 7SK snRNP complex that sequesters the positive transcription elongation factor (P-TEFb) and inhibits its activity (Quaresma et al., 2016; Nguyen et al., 2001; Uchikawa et al., 2015; Yang et al., 2001; Yik et al., 2003). Among all components of the 7SK snRNP complex and its close interactors, only the 7SK transcript was affected by treatment with GTX (Fig. 4b). Consistent with our observations after treatment of CD4+ T cells obtained from aviremic participants (Supplementary Fig. 7c), we did not detect significant change in expression of NF-kB, oxidative stress, apoptosis and T-cell effector function related genes after GTX treatment (Supplementary Fig. 8), indicating that GTX [20 nM] does not influence these pathways after 4 or 24 hours of stimulation.

As P-TEFb is an essential co-factor for Tat mediated HIV-1 transcription elongation (Ott et al., 2011; Mousseau and Valente, 2017) we examined whether GTX treatment affects P-TEFb activity. Phosphorylation of serine 2 residues within C-terminal domain (CTD) of RNA Polymerase II is a prerequisite for activation of transcription elongation and is mediated by the kinase activity of CDK9, a component of P-TEFb (Peterlin and Price, 2006). To examine whether GTX treatment resulted in Serine 2 RNA Pol II phosphorylation, we treated resting CD4+ T cells with GTX, the CDK9 inhibitor flavopiridol (FPD), and PMA or αCD3/αCD28 as positive control (Fig. 4c, Supplementary Fig. 9a). As expected, FPD treatment abrogated RNA Pol II phosphorylation, while PMA stimulation led to strong Ser2 RNA Pol II phosphorylation. Importantly, treatment with GTX caused an increase in phosphorylation of the CDK9 target, RNA Polymerase II Serine 2 in three independent donors tested (Fig. 4c and Supplementary Fig. 9a). The fact that short term GTX treatment enhanced P-TEFb activity in resting CD4+ T cells without cellular activation (Fig. 2g, 3d and 4c), led us to hypothesize that GTX treatment may cause destabilization of the 7SK snRNP complex leading to the release of free (active) P-TEFb. Active P-TEFb would then become available for transcription elongation at the latent HIV-1 LTR, a critical step required for HIV-1 latency reversal (Jonkers and Lis, 2015; Yukl et al., 2018). To test this, we performed glycerol gradient sedimentation experiments after treatment of resting CD4+ T cells with GTX (Figs. 5a-b, Supplementary Fig 9b). Indeed, 20 nM GTX treatment of CD4+ T cells for 4 hours resulted in release of free P-TEFb, from its inhibitory higher molecular weight 7SK snRNP complex, as shown by western blot analysis depicting distribution of the P-TEFb component CDK9 over high and low molecular weight fractions (Fig. 5a and Supplementary Fig 9b). As expected, control treatment of CD4+ T cell lysates with RNase A, which digests the 7SK RNA scaffold resulted in disassembly of the 7SK snRNP complex and subsequent release of free P-TEFb, which eluted at lower molecular weight fractions (Figs. 5a-b, Supplementary Fig. 9b). Quantification of free versus 7SK snRNP-sequestered P-TEFb demonstrates a significant GTX-induced release of free P-TEFb from the 7SK snRNP complex in CD4+ T cells from three independent donors (Fig 5b). To understand better which component of the 7SK snRNP complex may be targeted by GTX, we performed *in silico* docking experiments employing two independent software packages Chimera and Achilles. We modelled GTX against all essential components of the complex separately, as complete crystal structure of 7SK snRNP is not yet available. Strikingly, we observed preferential binding of GTX into the hydrophobic pocket of LARP7, which in physiological conditions is responsible for binding stem loop 4 (SL4) of the 7SK RNA (Fig. 5c-d and Supplementary Fig. 10). Interestingly, LARP7 is a critical component, responsible for stabilization of the 7SK snRNP complex as its depletion has been shown to lead to decreased levels of 7SK RNA with concomitant increase in free P-TEFb levels (Krueger et al., 2008). Moreover, consistent with our data, depletion of 7SK RNA was shown to result in release of free P-TEFb while depletion of LARP7, and to a lesser extent that of another 7SK snRNP complex structural component MEPCE, caused increased Tat-mediated transactivation of the HIV-1 LTR (Krueger et al., 2008). Our experimental data together with this modelling exercise and previously published data is consistent with a model in which GTX interferes with the binding of SL4 of 7SK RNA into the hydrophobic pocket of LARP7, which results in destabilization of the complex and subsequent release of P-TEFb and 7SK RNA (Fig. 5e). In resting CD4+ T cells, 7SK RNA is then degraded, and free P-TEFb made available for recruitment to the paused RNA Polymerase II at the latent HIV LTR by the Tat-TAR axis. CDK9 then phosphorylates CTD of RNA Polymerase II, leading to activation of proviral transcription elongation (Fig. 5e).

## Discussion

Distinct classes of LRAs have been shown to target different subpopulations of proviruses (Abner et al., 2018; Battivelli et al., 2018; Chen et al., 2017; Stoszko et al., 2019). Thus far none of the clinically tested LRAs has been able to induce strong viral expression or to significantly deplete the latent reservoir in patients, pointing to potential limitations of single treatments (Spivak and Planelles 2017; Rasmussen and Søgaard 2018). The heterogeneous nature of latent HIV integrations, the complex molecular mechanisms that contribute to maintaining HIV latency, together with individual genetic variability, dictate that a cocktail of stimulatory compounds targeting distinct cellular and HIV gene regulatory pathways would be most effective to activate the latent reservoir in HIV-1 infected patients (Stoszko et al., 2019). Indeed, preclinical studies demonstrate that combination treatments can result in synergism and lead to stronger HIV-1 latency reversal (Bouchat et al., 2016; Stoszko et al, 2015; Marian et al., 2018; Hashemi et al., 2018; Darcis et al., 2015; Abner and Jordan, 2019; Laird et al., 2015; Rasmussen and Lewin, 2016; Stoszko et al., 2019). Here we found that GTX, a molecule that targets the HIV-1 transcription elongation step, strongly synergized with LRAs that de-repress chromatin structure, namely BAF and HDAC inhibitors, which respectively target complexes that position the latent LTR chromatin in a repressive configuration (Rafati et al., 2011), or deacetylate histones compacting LTR chromatin. Interestingly, co-treatment of primary latently infected CD4+ T cells with GTX and BET inhibitors resulted in additive and not synergistic increase in HIV transcription. This observation is consistent with the fact that these compounds target the same mechanistic step, transcription elongation at the HIV-1 LTR. Our data highlights the attractiveness of simultaneous pharmacological targeting of mechanistically distinct steps in HIV-1 transcription regulation, namely at the level of chromatin structure, transcription factor-induced activation, as well as transcription elongation. We postulate that more robust latency reversal will be observed when compounds targeting these three mechanistic steps are combined, and further optimized by interventions that address blocks occurring post-transcriptionally, at the polyadenylation and splicing stages, which may hinder translation of HIV-1 genes (Yukl et al., 2018). In addition, consistent with the importance of targeting transcription elongation, shown recently to be a major rate limiting step in HIV-1 latency reversal in patient CD4+ T cells (Yukl et al., 2018), *ex vivo* GTX treatment of CD4+ T cells obtained from c-ART suppressed aviremic HIV-1 infected patients demonstrated significant increase in levels of cell associated HIV-1 RNA in all patients examined.

To our knowledge GTX is the first molecule described to cause direct disruption of the 7SK snRNP complex with subsequent release of active P-TEFb. Given the significance of P-TEFb in HIV transcription elongation and the current lack of available molecules able to mediate its release from the inhibitory 7SK snRNP complex, GTX may be a promising candidate, not only in context of HIV-1 latency reversal, but also in other diseases in which P-TEFb plays a prominent regulatory role (Kohoutek, 2009). Our modeling exercises provide insight towards a mechanism where GTX competes with 7SK RNA for the hydrophobic pocket of LARP7, causing destabilization and disassembly of the 7SK snRNP complex leading to release of P-TEFb and subsequent degradation of the 7SK RNA scaffold as observed in our experiments (Fig. 4, 5 and Supplementary Figs. 9 and 10). Indeed LARP7 has been shown to be a critical scaffold required for the stability of the 7SK snRNP complex; consistent with our observations, RNA interference mediated depletion of LARP7 resulted in elevated levels of free P-TEFb and increased Tat-mediated HIV-1 expression, concomitant with degradation of 7SK RNA (Krueger et al., 2008). To our knowledge, GTX is the only compound known to directly target 7sk snRNP for disruption.

Thus far gliotoxin has been regarded as a toxin and a virulence factor of *Aspergillus* fungi, shown to function as an immunosuppressant that inhibits phagocytosis, blocks Nf-kB signaling and cytokine production, and induce ROS formation (Choi et al., 2007; Kwon-Chung and Sugui, 2009; Sakamoto et al., 2015; Scharf et al., 2016; Stanzani et al., 2005; Wichmann et al., 2002; Yamada et al., 2000). These properties are ascribed to the molecule’s unusual disulfide bridge, that may mediate the activity of GTX by cycling between a reduced and oxidized state (Scharf et al., 2016). However, it is important to note that all of the above mentioned effects of GTX were observed at high concentrations, in the 100nM to 5 μM range, concentrations at which we also observed significant toxicity and cell death (Fig. 2, Supplementary Figs. 3, 4, 6). Interestingly, serum concentrations of GTX reported in patients with aspergillosis are found to be between 200 nM and 2.4 μM (Lewis et al., 2005), more than 10-100 orders of magnitude higher than concentrations we have shown to effectively reverse HIV-1 latency in primary T cells and cells of c-ART suppressed HIV-1 infected participants (Fig. 3-4). Taken together, these observations suggest that lower GTX concentrations in the 20nM range, at which effective HIV-1 latency reversal is observed in primary CD4+ T cells without associated toxicity, global T cell activation, or interference with capacity for CD8+ T cells activation, will be physiologically achievable in a therapeutic context. Our data therefore supports the potential of GTX for inclusion in combination latency reversal therapeutic approaches in a safe concentration range. Finally, our data underlines the power of coupling a screen of fungal extrolites, which comprise largely unexplored plethora of bioactive chemical entities, with bioassays and state of the art fractionation and mass spectrometry/NMR as a strategy to identify and characterize novel compounds with therapeutic potential.

## Materials and Methods

### Cells and culture conditions

Jurkat latent (J-Lat) cell lines A2 and 11.1 were cultured in RPMI-1640 media supplemented with 10% FBS and 100 µg/ml penicillin-streptomycin at 37 °C in a humidified 95%-air-5%CO_2_ incubator. Primary CD4+ T cells were cultured in RPMI-1640 media supplemented with 7% FBS and 100 µg/ml penicillin-streptomycin at 37 °C in a humidified 95%-air-5%CO_2_ incubator.

### Preparation of fungal supernatants

We screened 115 species of filamentous fungi (Table S1, Fig. S1A). Species belonged to 28 orders (43 families) of the fungal Kingdom. The majority were of ascomycetous affinity including ascomycetous yeasts (Saccharomycetales: four species), 12 were of basidiomycetous affinity including three basidiomycetous yeasts (Trichosporonales, Tremellales), and three species of lower fungi (Mucorales)(Gostinčar et al., 2018). Rationale for choice was the expected production of a wide array of metabolites, which are known to be more pronounced in fungi living in habitats with environmental stress or microbial competition. Particularly versatile nutrient scavengers in Eurotiales and Hypocreales are established metabolite and toxin producers. Onygenales are cellulose and keratin degraders, and contain a large cluster of mammal pathogens with alternating environmental life cycles. Members of Capnodiales are saprobes in environments with stressful microclimates such as rock, glass and indoor. Identity of all strains was verified by rDNA ITS (Internal Transcribed Spacer) and partial nucLSU (D1/D2 of Large SubUnit) sequencing.

Strains were derived from lyophilized storage in the reference collection of the Centraalbureau voor Schimmelcultures (housed at Westerdijk Fungal Biodiversity Institute, Utrecht, The Netherlands). After opening, contents of vials were taken up in 1 mL malt peptone and distributed on Malt Extract Agar (Oxoid) culture plates. Strains were maintained MEA slants and subcultured regularly until use. Mycelial fragments of one-week-old colonies grown on MEA at 25°C were used as inocula for 500-mL flasks containing 150 mL RPMI (with glutamine) shaken at 180 r.p.m. and incubated at 25°C for 7 days, with one negative control. Flasks were checked daily until biomass reached 1/3 to 1/4 of the volume, then harvested by centrifugation at 14,000 *g* and filtered using 0.2 μm metal filters. Supernatants were transferred to falcon tubes and used for latency screens. One strain per genus was used in a first round; additional isolates, some of which were close relatives and others located at larger phylogenetic distance, were tested in case of positive response. Positive tests of strain *Aspergillus fumigatus* CBS 113.26 was thus followed by *A. fumigatus* CBS 154.89, CBS 117884, CBS 100074, *A. lentulus* CBS 117884, and *A. flavus* CBS 625.66.

### Fractionation and mass spectrometry characterization of active components

Complex fungal supernatants (3.3 mL) were twice extracted with ethylacetate (3 mL) in glass tubes (5 minute vortex) and centrifuged at 3000 rpm. Water phase was discarded and organic extract dried in vacuum concentrator. Lipids were removed by consequent hexane/water extraction (50/50, v/v, 4 mL total volume), aqueous phase was collected and dried. The dried material was reconstituted in a 50% methanol and such a solution loaded onto HLB, MCX and MAX cartridges (Fig. 2A-B) obtained from Waters Corporation (Prague, Czech Republic). The adsorbed compounds were desalted and stepwise eluted with increasing (5, 45 and 95%) organic solvent (MeOH) gradient (1% HCOOH) providing A, B and C sample variants, respectively (Fig. 2B). HLB was based on N-vinylpyrrolidone – divinylbenzene copolymer and provided 5-fractions. MCX was cation exchange sorbent represented the HLB material modified with SO_3_H^-^ groups (6-fractions). The best performance was provided anion exchange MAX cartridge providing 7-fractions. After each round of fractionation all samples were tested in J-Lat/S-Lat latency models, followed by quantitation GFP/luciferase expression and identification of fraction retaining activating component. As expected, originally less active fractions have become positive as the active components became populated during the enrichment process. The same phenomenon was observed when working with less or more active *Aspergillus* strains (Fig. S2B).

The most active 7B/7C fractions were further fractioned on HLB cartridge (11-samples) and components dereplicated by Cyclobranch software (Novák et al., 2017). Compound matching against the annotated database of *Aspergillus* secondary metabolites has revealed a set candidate compounds further selected for latency reversal testing. Gliotoxin was present in active fractions (Figs. 2B and S2B).

Detailed examination of commercial (GTX, Cayman and ApexBio) and natural gliotoxins was performed by high performance liquid chromatography (HPLC) and Fourier transform ion cyclotron resonance (FTICR) mass spectrometry. SolariX XR 12T FTICR instrument (Bruker Daltonics, Billerica, USA) was operated in positive ion mode with electrospray ionization. Separation of GTX components was performed on Acquity UPLC HSS T3 analytical column (Waters, Prague, Czech Republic) with 1.0×150 mm dimensions and 1.8 um particle size. The analysis was carried out at 30 °C and 30 uL/min flow rate with the following A/B gradient: 0 min – 30% B; 30 min - 95% B; 40 min - 95% B; 50 min - 30% B; 60 min - 30% B. The gradient components A and B were represented by 0.1% formic acid in water or acetonitrile, respectively. One minute time windows around S2 and S3/S4 gliotoxins were used for fraction isolation. The preparative chromatography was performed both with commercial gliotoxin as well as with *A. fumigatus* 100074 strain.

### Flow cytometry and annexin V staining

GFP expression of cell lines J-Lat A2 and 11.1 and phenotype of spin infected primary CD4+ T cells at the moment of reactivation were analyzed by FACS (fluorescence-activated cell sorting) as described previously (Stoszko et al. 2015). For Annexin V staining 10^6^ cells were washed with PBS supplemented with 3% FBS and 2.5mM CaCl_2_ and stained with Annexin V-PE (Becton and Dickinson) for 20 min at 4 °C in dark in the presence of 2.5mM CaCl_2_. Stained cells were washed with PBS/FBS/CaCl_2_, fixed with 1% formaldehyde and analyzed by FACS.

### Primary CD4+ T cell isolation and infection

Primary CD4+ T cells were isolated from buffy coats from healthy donors by Ficoll gradient (GE Healthcare) followed by negative selection with RosetteSep Human CD4+ T Cell Enrichment Cocktail (StemCells Technologies). Isolated cells were left overnight for recovery and used for spin-infection according to Lassen and Greene method as described previously (Spina et al. 2013, Stoszko et al. 2015). Briefly, CD4+T cells were infected with the PNL4.3.LUC.R-E-virus by spinoculation (120 min at 1200 g) and cultured for three days in RPMI 10% FBS and 100 μg/ml penicillin-streptomycin in presence of Saquinavir Mesylate (5 μM). Three days after infection cells were treated as indicated in presence of Raltegravir (30 μM). Cells were harvested 24 h after treatment and luciferase activity was measured using Luciferase Assay System (Promega, Leiden, The Netherlands). Infections were performed using pseudotyped particles obtained by co-transfecting HXB2 Env with pNL4.3.Luc.R-E-plasmid into HEK 293T cells using PEI, 48 and 72 h post transfection, supernatants containing pseudotyped virus were collected, filtered with 0.45µm filter and stored at –80 °C. Molecular clones pNL4.3.Luc.R-E- and HIV-1 HXB2-Env, and antiretroviral drugs Saquinavir Mesylate and Raltegravir were kindly provided by the Centre for AIDS Reagents, NIBSC. HIV-1 molecular clone pNL4.3.Luc.R-E- and HIV-1 HXB2-Env expression vector were donated by Dr. Nathaniel Landau and Drs Kathleen Page and Dan Littman, respectively.

### HIV-1 latency reversal in primary CD4+ T cells isolated from aviremic patients

Primary CD4+ T cells were isolated as described previously (Stoszko et al., 2015). Three million CD4+ T cells were plated in triplicate at the cell density of 10^6^/ml and treated as indicated. After 24 hours cells were lysed and total RNA was isolated as described below. Written informed consent was obtained from all patients involved in the study. The study was conducted in accordance with ethical principles of the Declaration of Helsinki. The study protocol and any amendment were approved by The Netherlands Medical Ethics Committee (MEC-2012-583).

### Total RNA isolation and quantitative RT-PCR

Cells were lysed in TRI reagent and RNA was isolated with Total RNA Zol-out kit (A&A Biotechnology), cDNA synthesis and qPCR was performed as described previously (Stoszko et al., 2015). Gene expression was calculated using 2^-ΔΔCt^ method (Schmittgen and Livak, 2008), expression of GAPDH was used for normalization. Absolute quantification of cell-associated *pol* RNA was performed as described previously (Stoszko et al., 2015). Briefly, qPCR was performed in a final volume of 25 μl using 4 μl of cDNA, 2.5 μl of 10× PCR buffer (Life Technologies), 1.75 μl of 50mM MgCl_2_ (Life Technologies), 1 μl of 10 mM dNTPs (Life Technologies), 0.125 μl of 100 μM Pol Forward primer (HXB2 genome 4901 → 4924), 0.125 of 100 μM Pol Reverse primer (HXB2 genome 5060 → 5040), 0.075 μl of 50 μM of Pol Probe, and 0.2 μl Platinum Taq polymerase (Life Technologies). The lower limit of detection of this method was of 20 copies of HIV-1 RNA in1 μg of total RNA. The absolute number of polcopies in PCR was calculated using a standard curves ranging from 7 to 480 copies of a plasmid containing the full-length HIV-1 genome (pNL4.3.Luc.R-E-). The amount of HIV-1 cellular associated RNA was expressed as number of copies/μg of input RNA in reverse transcription. Preparations of cell-associated RNA were tested for potential contamination with HIV-1 DNA and-or host DNA by performing the PCR amplification in the presence and absence of reverse transcriptase.

Primers used for RT-qPCR:

pol forward 5’GGTTTATTACAGGGACAGCAGAGA3’

pol reverse 5’ACCTGCCATCTGTTTTCCATA3’

pol probe [6FAM] AAA ATT CGG TTA AGG CCA GGG GGA AAG AA[BHQ1]

TNFα forward 5’TAGGCTGTTCCCATGTAGCC3’

TNFα reverse 5’CAGAGGCTCAGCAATGAGTG 3’

IL-2 forward 5’ACCTCAACTCCTGCCACAAT 3’

IL-2 reverse 5’GCACTTCCTCCAGAGGTTTG 3’

INFγ forward 5’ATAATGCAGAGCCAAATTGTCTCC3’

INFγ reverse 5’ATGTCTTCCTTGATGGTCTCCAC 3’

CD25 forward 5’ATCAGTGCGTCCAGGGATAC 3’

CD25 reverse 5’GACGAGGCAGGAAGTCTCAC 3’

BAK1 forward 5’GGTTTTCCGCAGCTACGTTTTT 3’

BAK1 reverse 5’GCAGAGGTAAGGTGACCATCTC3’

BAX forward 5’CCCGAGAGGTCTTTTTCCGAG 3’

BAX reverse 5’CCAGCCCATGATGGTTCTGAT3’

BCL2 forward 5’GGTGGGGTCATGTGTGTGG3’

BCL2 reverse 5’CGGTTCAGGTACTCAGTCATCC3’

CASP-3 forward 5’CATGGAAGCGAATCAATGGACT3’

CASP-3 reverse 5’CTGTACCAGACCGAGATGTCA3’

AKT1 forward 5’AGCGACGTGGCTATTGTGAAG3’

AKT1 reverse 5’GCCATCATTCTTGAGGAGGAAGT3’

NRF2 forward 5’TTCCCGGTCACATCGAGAG3’

NRF2 reverse 5’TCCTGTTGCATACCGTCTAAATC3’

FOXO3a forward 5’CGGACAAACGGCTCACTCT3’

FOXO3a reverse 5’GGACCCGCATGAATCGACTAT3’

KEAP1 forward 5’GCAATGAACACCATCCGAAGC3’

KEAP1 reverse 5’ACCATCATAGCCTCCAAGGAC3’

### Glycerol gradient sedimentation (Fig. S8)

Glycerol gradients were prepared as described previously (Contreras et al., 2007). Briefly, around 40 x 10^6^ primary CD4+ T cells isolated from healthy donors were either untreated or treated with GTX [20 nM] for 4 h, for RNAse A treatment. Cells were lysed for 30 min in buffer A supplemented with either 100U/ml RNasin (Promega) for untreated and GTX conditions, or with 100 µg/ml RNase A. RNase supplemented lysates were incubated for 10 min at room temperature to ensure efficient degradation of RNA. Lysates were fractionated by centrifugation in a SW41 Ti rotor at 38000rpm for 20h. 1mL fractions were collected in 2 ml tubes and subjected to Trichloroacetic Acid precipitation of proteins as described previously (Link and LaBaer, 2011). Briefly, to each 1 ml fraction 110 µl of ice-cold 100% trichloroacetic acid (TCA) was added and incubated on ice for 10 min. Then, 500 µl of ice-cold 10% TCA was added to each sample and incubated on ice for 20 min, followed by centrifugation at 20 000g for 30min. Supernatants were carefully removed and precipitates were gently washed with 500 µl of ice-cold acetone followed by centrifugation at 20 000g for 10 min. Supernatants were gently removed and dried at room temperature for about 10 min. Protein precipitates were then re-suspended in 50 µL of Laemmli loading buffer and subjected to 10% SDS-PAGE separation and detection of CDK9 (C-20, sc-484, Santa Cruz Biotechnology).

### RNA Polymerase II phosphorylation

Ten million primary CD4+ T cells were either untreated or treated with GTX, OTX-015 [1 µM], flavopiridol [500 nM], PMA [10 nM] or αCD3/CD28 beads (cell: bead ratio 1:1) for 6 h. Cell were lysed for 30min on ice with IP buffer (Stoszko et al., 2015) and subjected to western blot analysis using following antibodies: total RNAPII (N-20, sc-899 Santa Cruz Biotechnology), phospho-Ser2 RNAPII (H5, ab-24758, Abcam), CDK9 (C-20, sc-484, Santa Cruz Biotechnology), α-tubulin (ab6160, Abcam).

### Histone acetylation

Ten million primary CD4+ T cells were treated with gliotoxin concentration gradient, SAHA or left untreated for 4 h and then washed twice in PBS. Cells were lysed for 10 min at 4 °C in TEB buffer (PBS, 0.5 % triton X-100 (v/v), 2 mM phenylomethylsulfonyl fluoride (PMSF), 0.02 % (w/v) NaN_3_) at a density of 10^7^ cells per 1 ml of the buffer. Samples were centrifuged at 425 g for 10 min at 4 °C. Supernatants were discarded and cell pellets were washed in half the volume of TEB buffer used for lysis and centrifuged as before. Supernatants were discarded and pellets were resuspended in 0.2N HCl at a density of 4 x 10^7^ cells per ml. Histones were extracted overnight at 4 °C and then centrifuged at 425 g for 10 min at 4 °C. Supernatants were collected, protein concentration was determined by the Bradford assay and samples were subjected to SDS-PAGE western blot. Following antibodies were used in western blot analysis: anti-acetyl-histone H4 (06-598, Milipore) and anti-histone H2B (ab52484, Abcam).

### RNA sequencing and data analysis

Ten million primary CD4+ T cells were stimulated with 20 nM gliotoxin or left unstimulated for 4 h. Experiment was performed in duplicate on cells isolated from two buffy coats from healthy donors as described above. RNA was isolated as described above. cDNA libraries were generated using Illumina TruSeq Stranded mRNA Library Prep kit (Illumina). The resulting DNA libraries were sequenced according to the Illumina TruSeq Rapid v2 protocol on an Illumina HiSeq2500 sequencer. Reads were generated of 50 base-pairs in length. Reads were mapped against the GRCh38 human reference using HiSat2 (version 2.0.4). We called gene expression values using htseq-count (version 0.6.1), using Ensembl transcript annotation.

Heat maps were generated using MORPHEUS (https://software.broadinstitute.org/morpheus/index.html).

### Toxicity assay

Peripheral blood mononuclear cells (PBMC) were isolated by density gradient centrifugation (Ficoll-Hypaque, GE Healthcare life sciences) from buffy coats from healthy donors (Sanquin Amsterdam) and either used immediately or frozen in freezing media (90% FBS/10% DMSO) and stored short term at −80°C. For cytotoxicity testing, cells were cultivated in culture media RPMI 1640 (Life Technologies) supplemented with 10% FBS, 2 mM L-glutamine, 100 U/ml penicillin, and 100g/ml streptomycin-sulfate at a concentration of 1×10^6^ cells/ml in 24-well plates (Thermo Scientific) that were either uncoated (unstimulated cells) or coated with anti-human CD3 (1µg/ml, clone UCHT1, no azide/low endotoxin, BD Bioscience) and anti-CD28 (10µg/ml, clone CD28.2, no azide/low endotoxin, BD Bioscience) monoclonal antibodies (stimulated cells). The following LRA were added at the concentrations indicated in figures: Gliotoxin (GTX, ApexBio), SAHA-Vorinostat (Selleck Chemicals), caffeic acid phenethyl ester (CAPE, MP Biomedicals), and Romidepsin (RMD, Sigma Aldrich). LRA at indicated concentrations were added to the cultures and cells were continuously exposed for 72 hours. Since GTX was reconstituted in acetonitrile (ACN), and all other LRA in dimethyl sulfoxide (DMSO), both solvents were added to the DMSO/ACN control culture (ACN 1:10000, DMSO 1:2500) to control for the effect these solvents may have on cell viability.

### Flow cytometric for cytotoxicity assay

To examine the effect the LRA have on immune cell subpopulations, cell viability, activation and proliferation was analyzed by flow cytometry. Surface antigens were detected by incubating 0.8-1.0×106 cells with pre-determined optimal concentrations of monoclonal antibody mixes in FACS wash (FW, Hanks buffered saline solution (HBSS, Life Technologies)), 3% fetal bovine serum (FBS, Life Technologies), 0.02% NaN3, 2.5 mM CaCl2) for 20 min at 4° C, washed one times with FW and fixed with 1% paraformaldehyde. To determine proliferation, cells were stained with 0.07 µM CellTrace Far Red Cell Proliferation dye according to manufacturer’s protocol (ThermoFisher Scientific) before cultivating for 72 hours with either unstimulated or stimulated conditions in the presence of LRA.

The following directly conjugated monoclonal anti-human antibodies were used to analyze CD8+ T cells (CD3+CD8+), CD4+ T cells (CD3+CD4+), B cells (CD3-CD19+), monocytes (CD14+) and NK cells (CD3-CD56+): CD3-BV421 (clone UCHT1), CD4-BV650 (SK3), CD8-BV786 (RPA-T8), CD14-PE-Cy7 (61D3, eBioscience), CD19-PerCP-Cy5.5(HIB19, eBioscience), CD56-PE-Cy5.5(CMSSB, eBioscience), Annexin V-PE, CD25-Super Bright 600 (BC96, eBioscience). All antibodies were purchased from BD Biosciences unless otherwise indicated. Between 2 - 4×10^5^ events were collected per sample within 24 hours after staining on a LSRFortessa (BD Biosciences, 4 lasers, 18 parameters) and analyzed using FlowJo software (version 9.7.4, Tree Star). Data are represented as frequency within a defined population. Drug-specific cell death was calculated using the following formula:

[(% Drug-induced cell death – % cell death in DMSO/ACN only)/(100 - % cell death in DMSO/ACN only)]*100

### Molecular docking

Molecular docking was used to predict the most likely binding mode of gliotoxin to LARP7 C-terminal domain. The crystal structure of human LARP7 C-terminal domain in complex with 7SK RNA stem loop 4 (PDB ID code 6D12) was optimized using PDB-REDO (Joosten et al., 2014) and used as a template to define the receptor for the docking simulation. The resulting pdb file was manually adapted for input into the docking procedure by elimination of protein chain B and RNA domain C and replacement of Selenium atoms (present as selenomethionin, incorporated for phasing purposes; (Eichhorn et al., 2018)) by sulfur. GTX ligand structure was built and energy minimized using the program Chimera (Pettersen et al., 2004). Molecular docking of GTX to LARP7 C-terminal domain was performed using Achilles Blind Docking server (https://bio-hpc.ucam.edu/achilles/) and Chimera’s AutoDock Vina function (Trott and Olson, 2010). The resulting solutions were ranked based on highest binding affinity (or lowest binding energy). Figures were created using PyMol software (Schrödinger, LLC, 2015).

### Quantification and statistical analysis

#### Western blot quantification

Differential band density was quantified by ImageQuant TL software using area and profile-based toolbox. For glycerol gradient western blot quantification an area frame was defined for all bands (total CDK9 protein content in all fractions), complex-bound CDK9 bands (heavy fractions) or free CDK9 bands (light fractions) were defined. The three area frames were measured for total density after background subtraction (local average). Relative complex-bound CDK9 or free CDK9 percentage was calculated regarding the untreated control (total CDK9 abundance). For immunoprecipitation western blot quantification an area for each LARP7 and CDK9 band was defined and total density was measured after background subtraction (local average). LARP7 abundance was first normalized to CDK9 abundance for each lane and relative abundance was calculated regarding untreated control. Statistical significance was calculated using ANOVA followed by Tukey’s test (p<0.01).

#### Statistical significance

Statistical significance was calculated as indicated in figure legends. Analyses were performed using Prism version 7.03 (GraphPad software).

#### Data and code availability

Sequencing data that support the findings of this study will be deposited in GEO and the accession codes will be provided upon request.

## Acknowledgement

TM received funding from the European Research Council (ERC) under the European Union’s Seventh Framework Programme (FP/2007-2013)/ERC STG 337116 Trxn-PURGE, the Dutch AIDS Fonds grant. 2014021 and Erasmus MC mRACE research grant.

Y.K was supported by the Technology Innovation Team Project of Guiyang [20161001]005; Guiyang Science and Technology Project [2017] No.5-19.

VH was supported by the Ministry of Education, Youth and Sports of the Czech Republic (LO 1509).

RP was supported by the Dutch AIDS Fonds grant. 2016014

JL is supported by the gravitation program CancerGenomiCs.nl from the Netherlands Organisation for Scientific Research (NWO). Supported in part by a grant awarded by Worldwide Cancer Research to PDK

## Author contribution

M.S., A.M.S.A, A.S., M.D.R. E.N. Y.M., M.J.N., R.C., J.K., R.P., P.B., A.B., TW.K., E.dC., R-J.P. M.Su. and C.R conducted the experiments; M.S., Y.M., R-J.P, J.H.G.L, A.V., P.D.K., V.H., S.de.H. and TM designed the experiments and wrote the paper.

The authors declare no competing interests.

## Supplementary Figures

**Supplementary Fig. 1.**
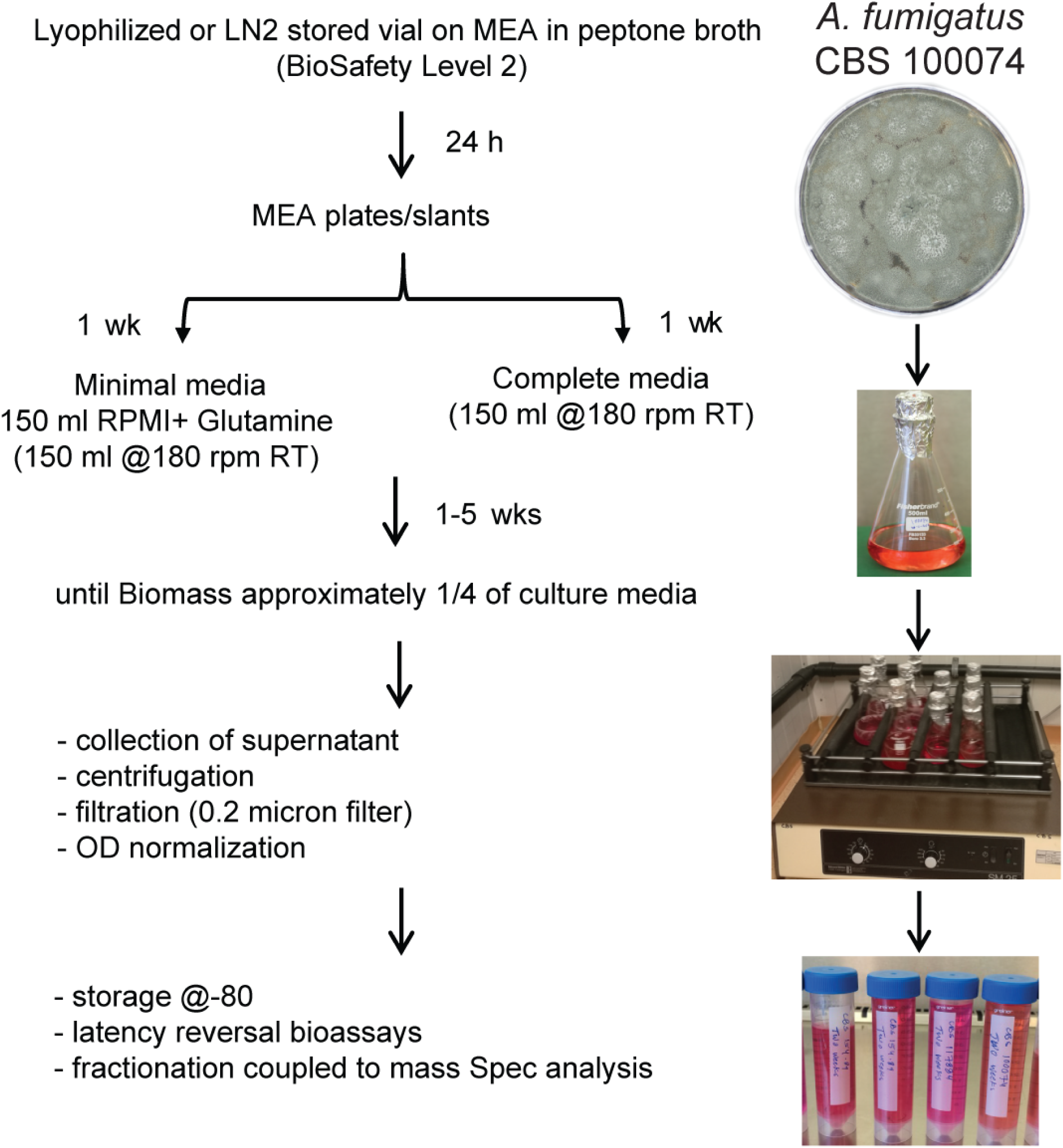
Schematic representation of the strategy used and details to culture fungi and prepare fungal growth supernatants for latency reversal bioassays.

**Supplementary Fig. 2.**
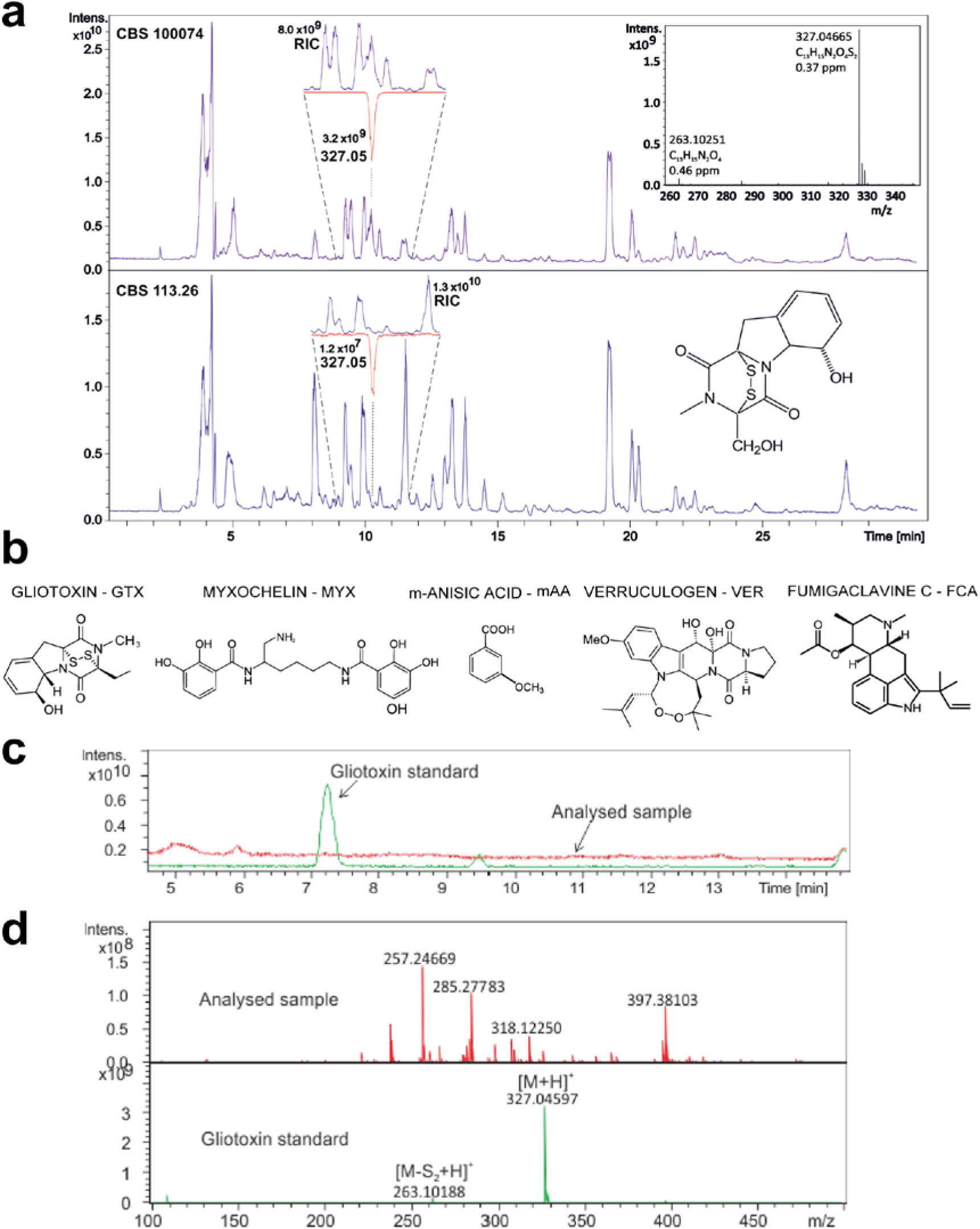
Identification of constituents of fungal supernatants. **a,** HPLC-MS separation of dichloromethane extracts of Aspergillus fumigatus (CBS 100074 and CBS 113.26) fermentation broths. The insets indicate different gliotoxin content representing here more th an 2.5 orders of magnitude (electrospray ionization). **b,** Commercially available, common constituen ts found in positive fractions that were tested for HTV-I latency reversal. **c-d,** Quest for GTX in CBS 625.66 supernatant. c, Total ion current from HPLC-MS of GTX standard (green curve) and analyzed sample (red curve). **d,** MS spectrum extracted from the chromatogram (7-8 min region).

**Supplementary Fig. 3.**
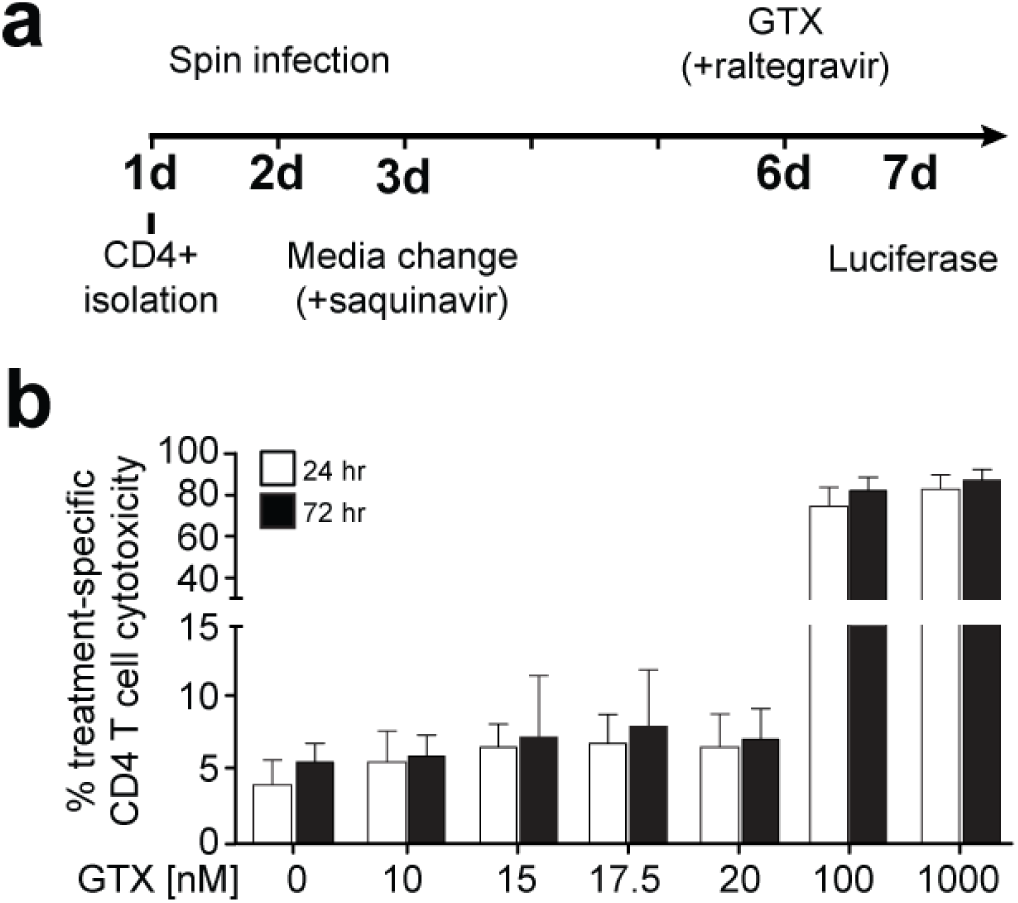
GTX treatment of primary latent HIV-1 infected CD4+ T cells at low concentrations does not induce cytotoxicity. **a,** Schematic of the strategy used to generate latent ex-vivo HIV-1 infected primary CD4+ T cells. **b,** Viability as shown by Annexin V staining of primary CD4+ T cells treated with increasing concentrations of GTX as indicated añer 24- and 72-hours treatment.

**Supplementary Fig. 4.**
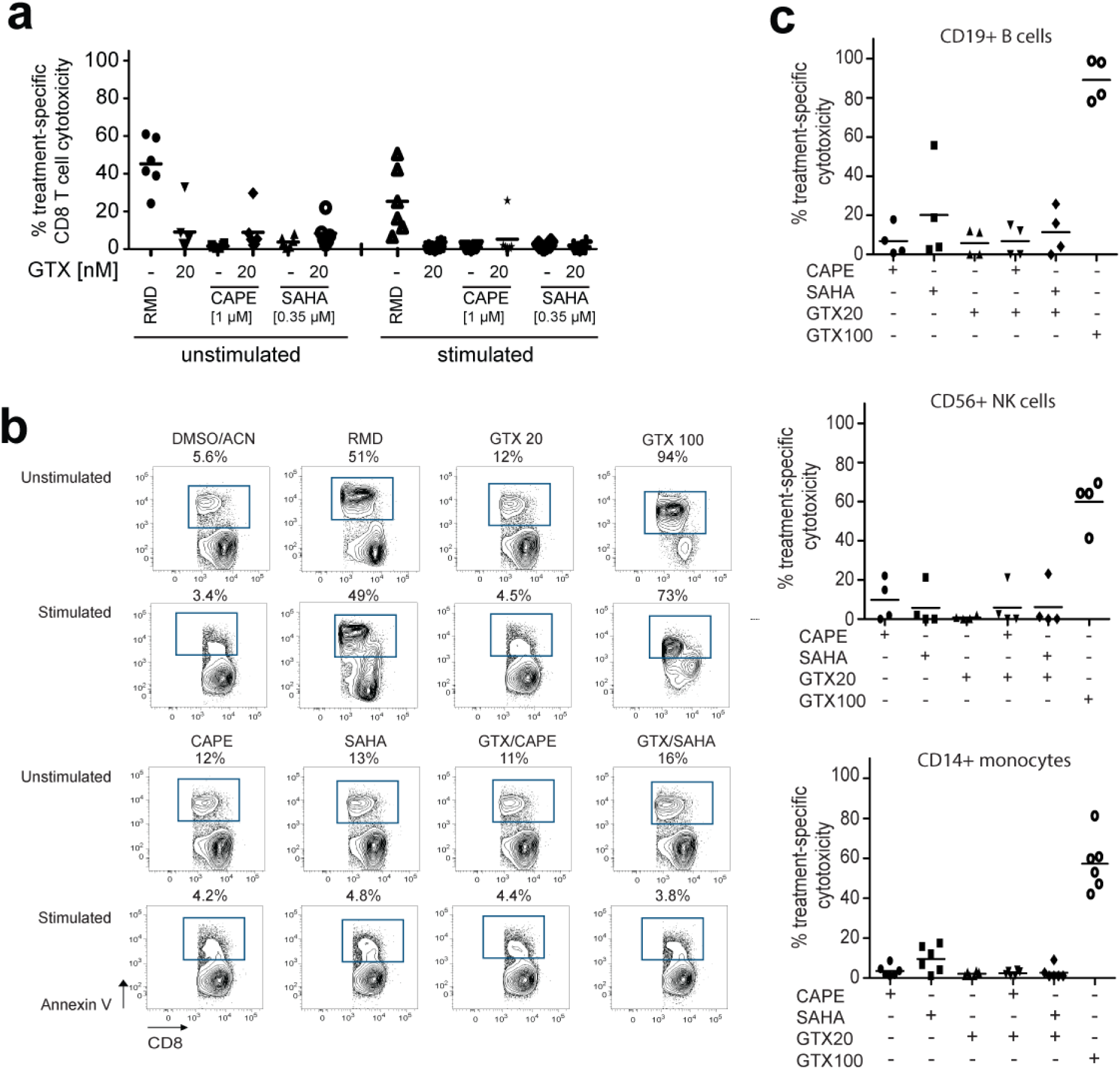
Materials and Methods Cells and culture conditions GTX [2 nM] treatment alone and eombined with the indicated LRAs is non-toxic in CD8+ T cells, B cells, NK cells and monoeytes. **a,** PBMC from a healthy donor were incubated with the indicated concentrations of either DMSO/ACN or LRA under unstimulated or plate-bound anti-CD3/anti-CD28 antibody stimulated conditions. After 72 hours exposure, cell death of CD8+CD3+ T cells was determined by Annexin V staining and flow cytometry. Each Symbol represents one healthy donor (n = 6 from 2-3 independent experiments), horizontal line depiets mean, **b,** Representative flow cytometry plots are shown for one healthy donor. Numbers in plot depict frequeney of gated Annexin V+ CD8+ T cells. **c,** GTX [20 nM] is non-cytotoxic for B cells (left panel), NK cells (middle panel) and monoeytes (right panel). PBMC from healthy donors were incubated with the indicated concentrations of LRA for 72 hours and the frequeney of drug-specific cell cytotoxicity was calculated for CD19+ B cells, CD56+ NK cells and CD14+ monoeytes.

**Supplementary Fig. 5.**
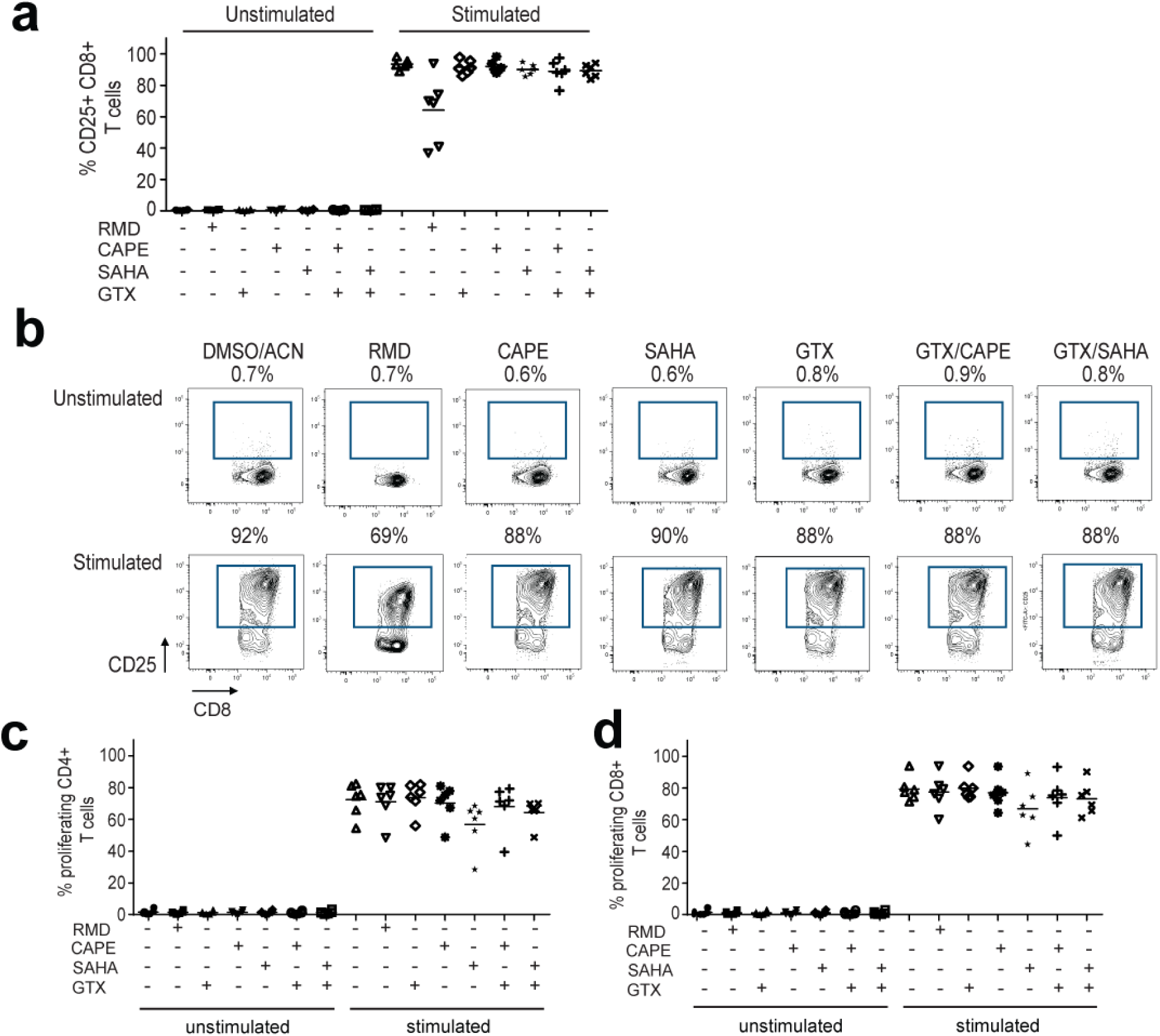
GTX [20 nM] does not induce T cell activation, not does it interfere with activation-mediatcd T cell proliferation. **a,** 20 nM GTX does not alter activation of CD8+ T cells. PBMC from healthy donors were incubated with the indicated LRAs for 72 hours either unstimulated or stimulated with aCD3/aCD28 antibodies. Panels depicts pooled data showing the ffequency of CD25+ cells within CD8+ T cells. Each symbol represents one healthy donor (n ***= 6*** from 2-3 independent experiments), horizontal line depicts mean, **b,** FACS panels represent the gating strategy used for analysis. **c,** CD4+ T cells and **d,** CD8+ T cells were treated as indicated and the frequency of proliferating cells was assessed by flow cytometry. Each symbol represents one healthy donor (n = 6 from 2-3 independent experiments), horizontal line depicts mean.

**Supplementary Fig. 6.**
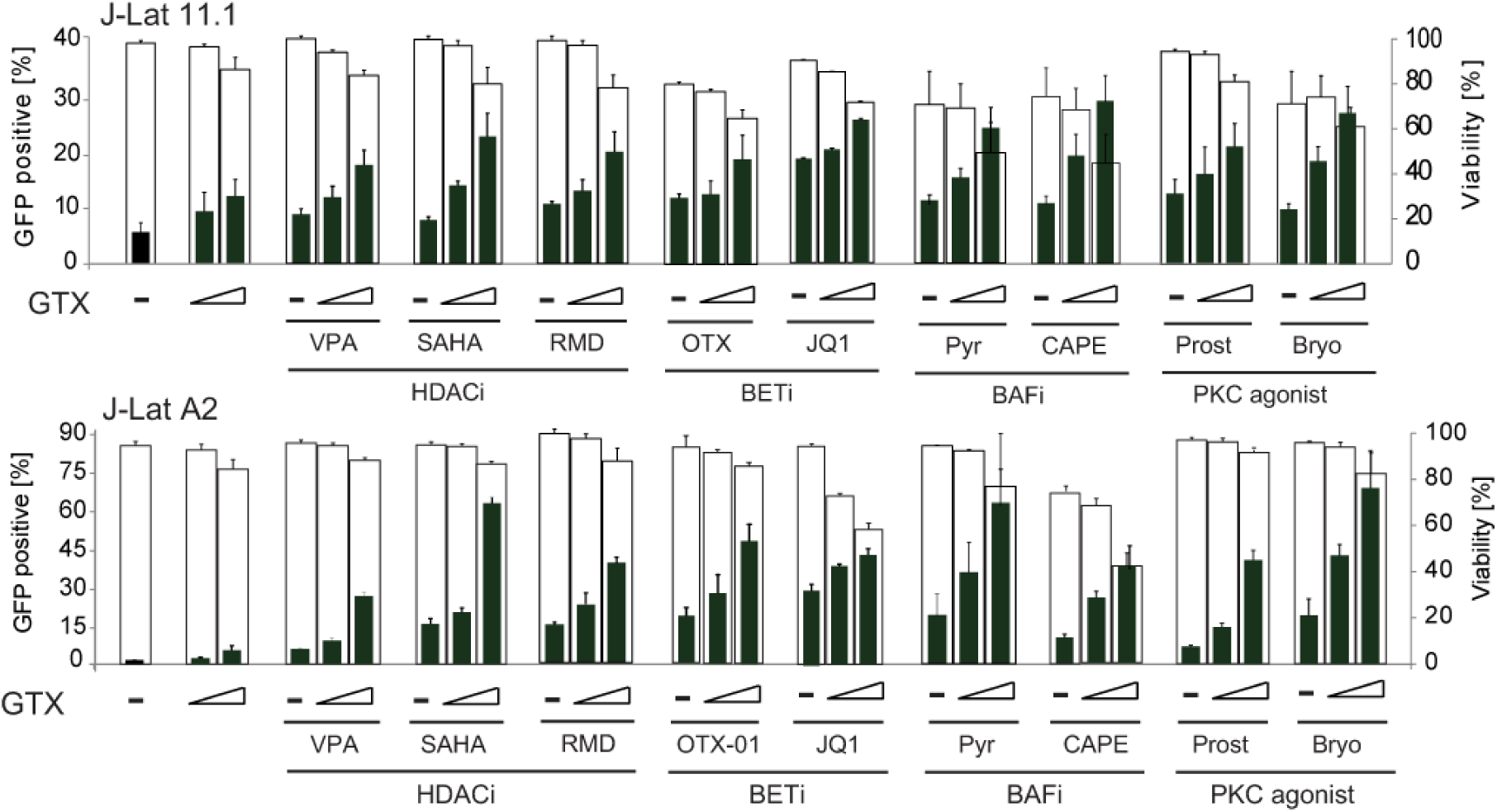
GTX enhances the lateney reversal activity of distinct LRA class molecules. J-Lat A2 or 11.1 cells were left untreated or treated with increasing GTX concentrations (0.25 pM and 0.5 pM) alone or in combination with other known LRAs as indicated for 48 hours. Lateney reversal (%GFP, left axis, green bars) and cell viability (%viable, right axis, empty bars) was then assessed by flow cytometry analysis. Data are presented as mean ± SD from three independen! experiments.

**Supplementary Fig. 7.**
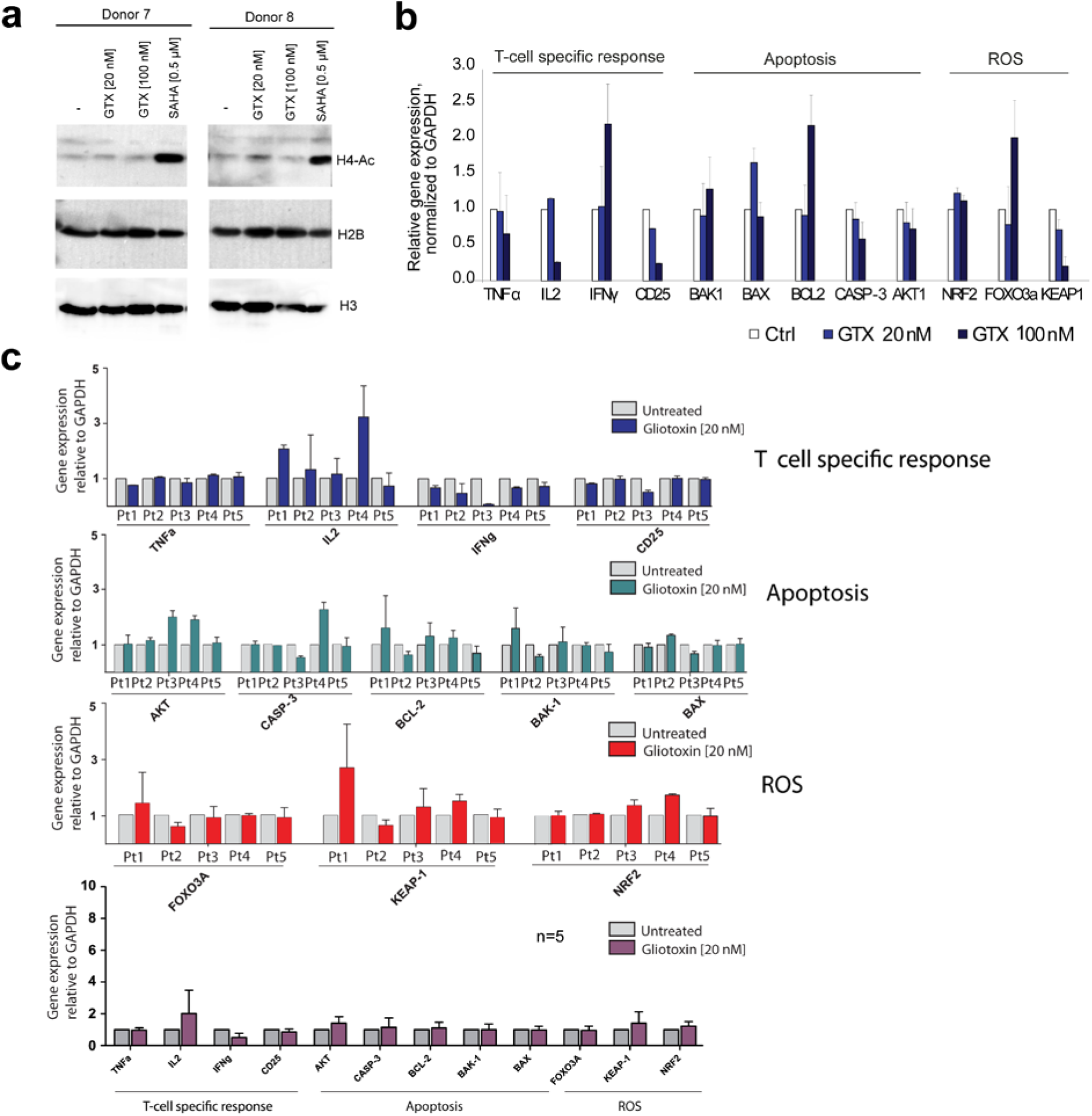
GTX [20-100 nM] is not an HDAC inhibitor and does not induce apoptosis,ROS or PKC target genes. **a,** GTX is not an HDAC inhibitor. Primary CD4+ T cells from two healthy blood donors were left untreated or treated with GTX or SAHA as indicated for 4 h and subjected to western blot analysis of acetylated histone H4, and total histones H2B and H3. b, GTX treatment of primary CD4+ T cells at lower concentrations does not affect expression of reactive oxygen species (ROS), apoptosis or T-cell specific response related genes. Cells were isolated from 2 healthy blood donors and treated with GTX or left untreated for4h as indicated. mRNA expression of T-cell effector functions, apoptosis and ROS related genes was assessed by real-time PCR. c, GTX treatment of CD4+ T cells obtained from aviremic participants does not induce ROS, apoptosis or T-cell specific responses. 20 nM GTX treatment of primary CD4+ T cells isolated from 5 aviremic patients for 24 hours does not affect expression of reactive oxygen species (ROS), apoptosis or T cell effector function related genes. mRNA expression of genes was assessed by real-time PCR, expression was normalized with GAPDH. Bottom panel represents averaged data from all 5 aviremic HIV+ participants.

**Supplementary Fig. 8.**
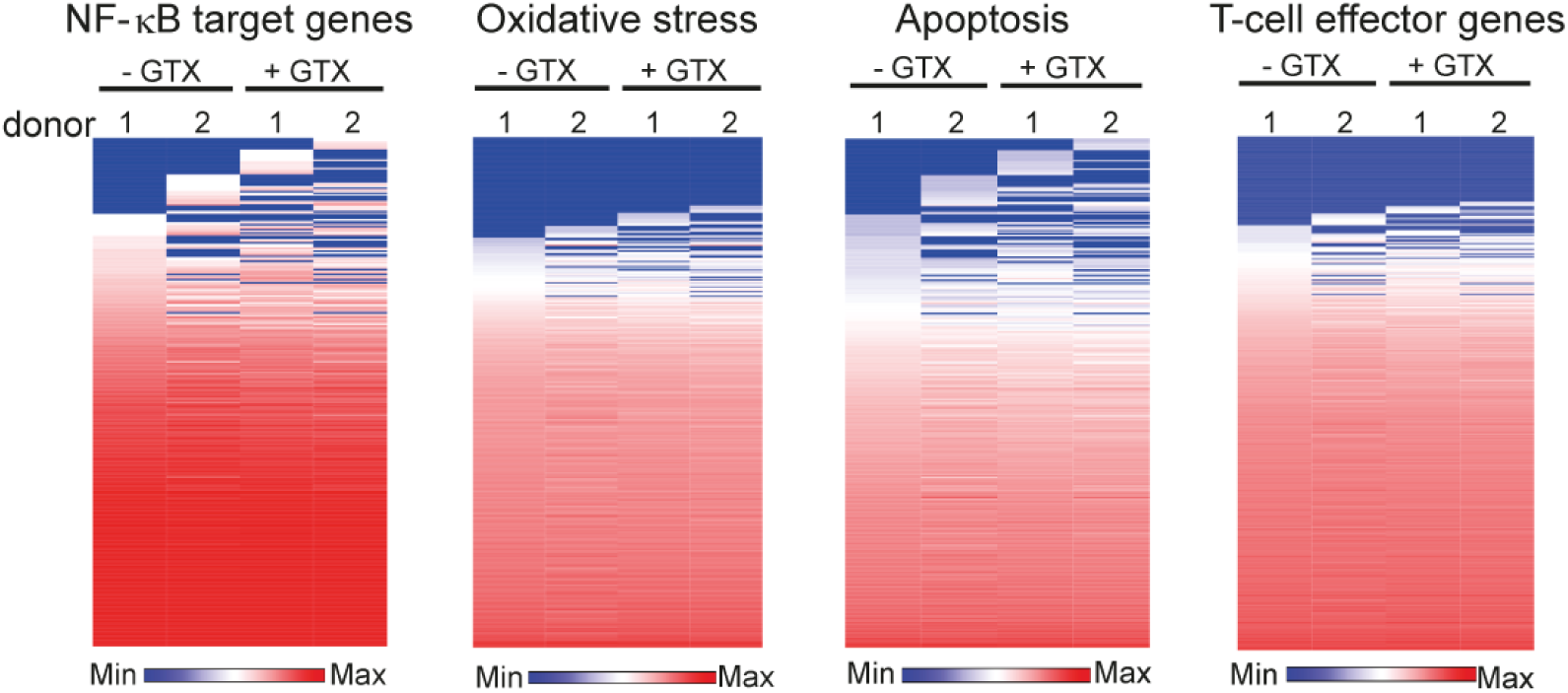
GTX [20 nM] treatment of resting CD4+ T cells does not influence the NF-kB, oxidative stress, apoptosis and T-cell effector pathways. Heatmaps of RNA-seq data for NF-0 B target genes, oxidative stress related genes, apoptosis related genes and T-cell effector genes demonstrates that 20 nM GTX does not influence these pathways. Primary CD4+ T cells from two healthy blood donors were left unstimulated or stimulated with GTX [20nM] for 4 hours, RNA was isolated and processed for RNA sequencing.

**Supplementary Fig. 9.**
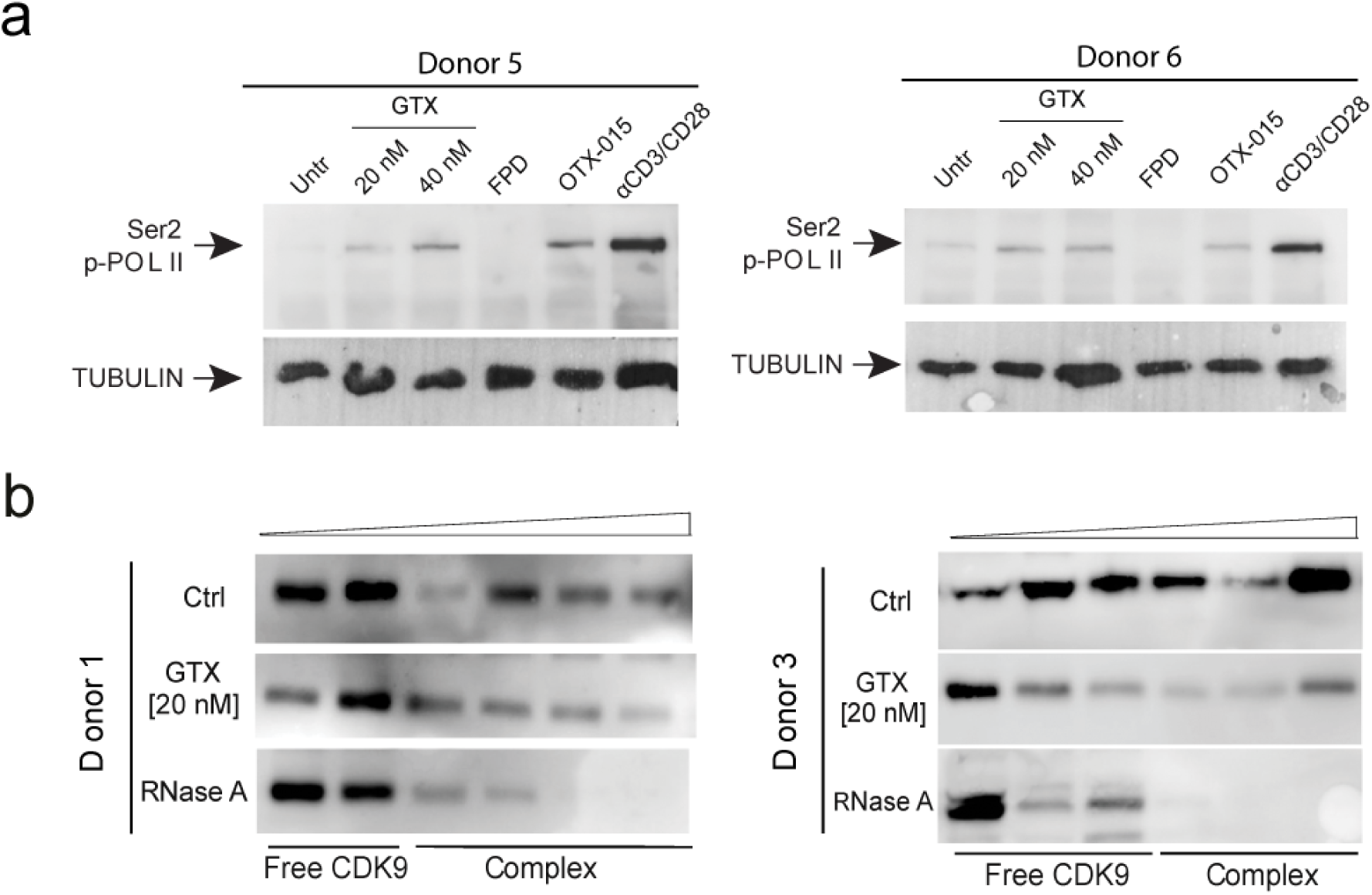
GTX treatment causes increased activity of P-TEFb by disrupting the inactive 7SK snRNP, leading to release of free P-TEFb. **a,** Phosphorylation of RNA polymerase II at serine 2 occurs in CD4+ T cells in response to treatment with GTX. Primary CD4+ T cells were untreated or treated for 6 hours with GTX [20-40 nM], BET inhibitor OTX-015 [1 pM], CDK9 catalytic inhibitor flavopiridol (FPD) [500 nM] and a-CD3/CD28 as a positive control. Western blot analysis was perfonned using antibodies specic for serine 2 phosphorylated RNA Pol II and tubulin as indicated. **b,** GTX treatment causes release of P-TEFb from sequestration within the inactive 7SK snRNP complex. Western blot analysis of glycerol gradient sedimentation of lysates from primary CD4+ T cell (from two additional donors) either untreated or treated with RNAse [100 pg/ml] or GTX [20nM] as indicated, using antibody specifically recognizing CDK9.

**Supplementary Fig. 10.**
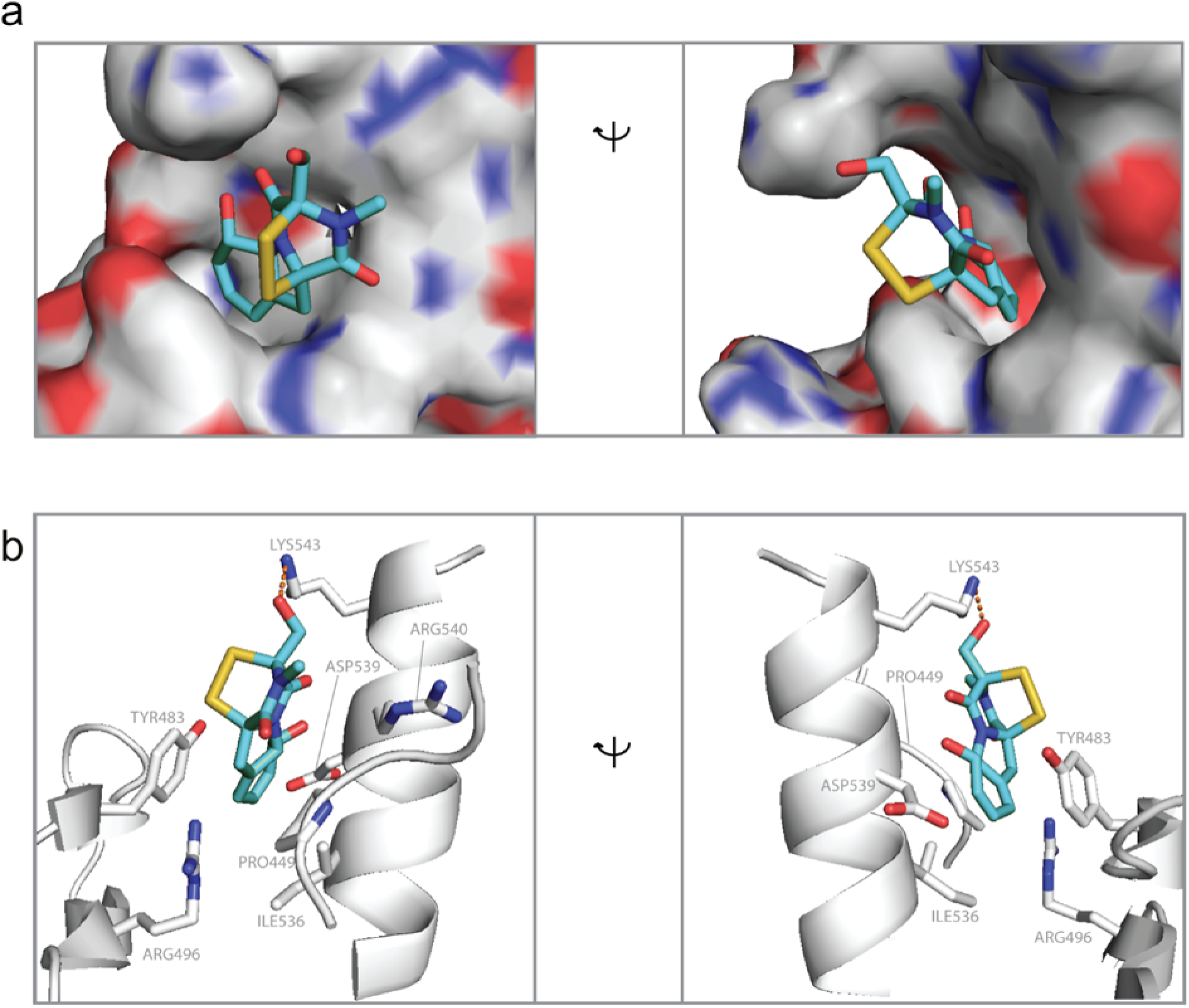
Molecular docking of GTX onto LARP7. **a,** Close up of hydrophobic pocket on the surface of LARP7 with bound GTX (highest scoring prediction from Achilles coordinates) in two orientations. LARP7 surface color code is red for oxygen, blue for nitrogen, grey for carbon (hydrophobic). Color code for GTX as in panel B. **b,** Details of interactions formed between LARP7 and GTX. Color code as in panel B. The predicted hydrogen bond between GTX and Lysine 543 from LARP7hydrophobic pocket on the surface of is indicated with a dashed line. Side chain atoms of Asp539, Ile536, Pro449 and Tyr483 make favorable van der Waals contacts with GTX. All images were generated with the use of PyMol software.

**Supplementary Table 1.**
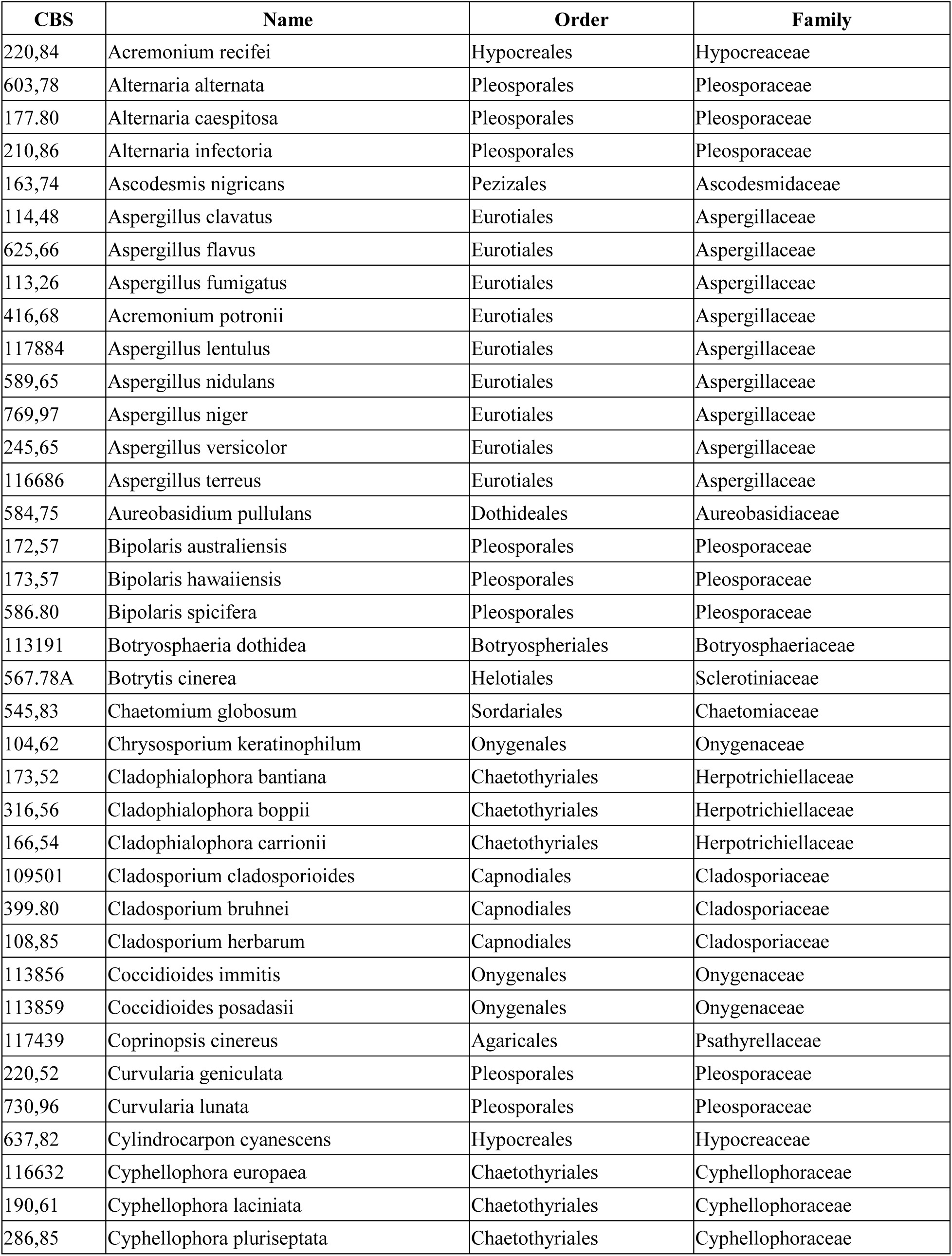
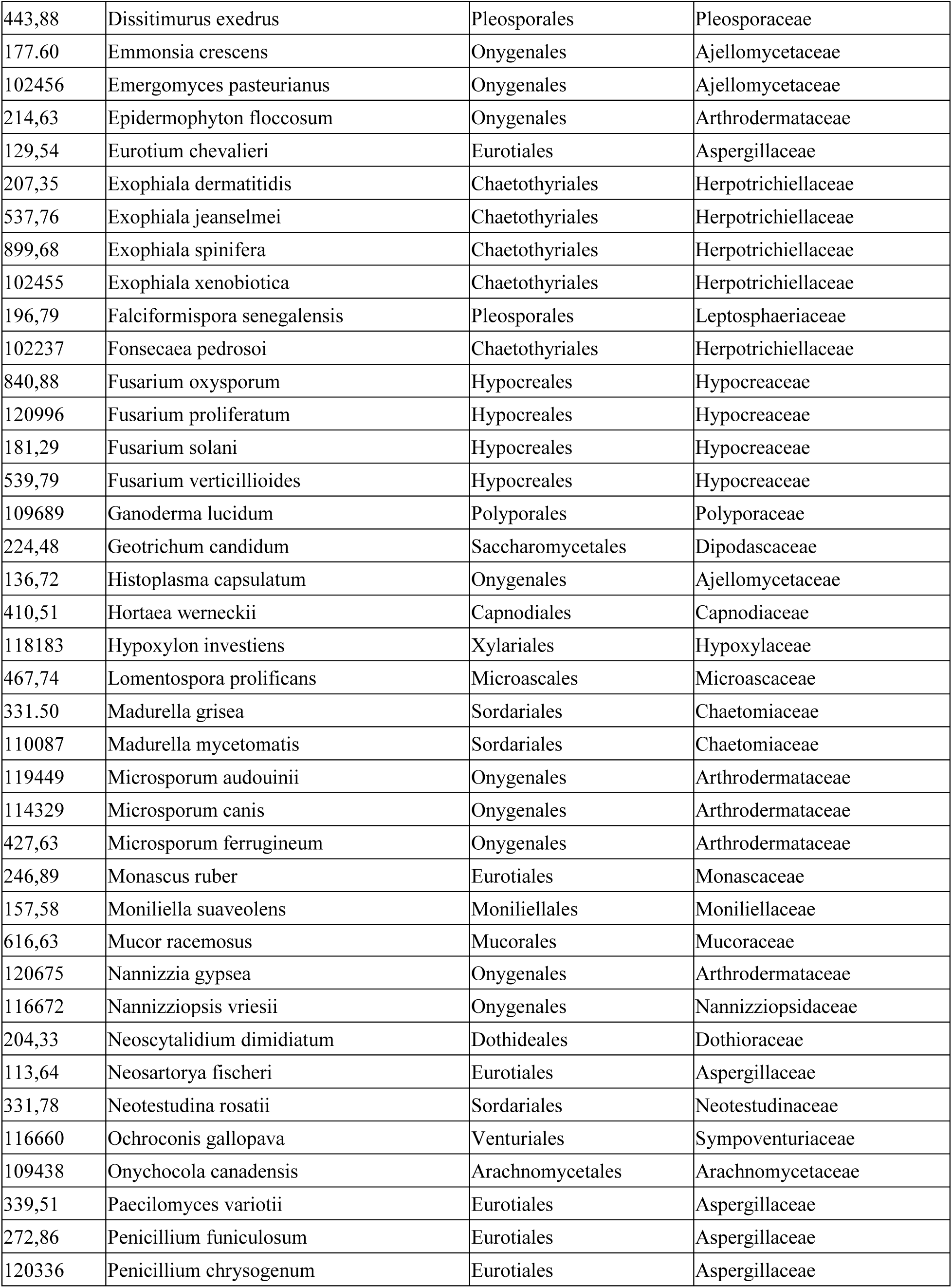
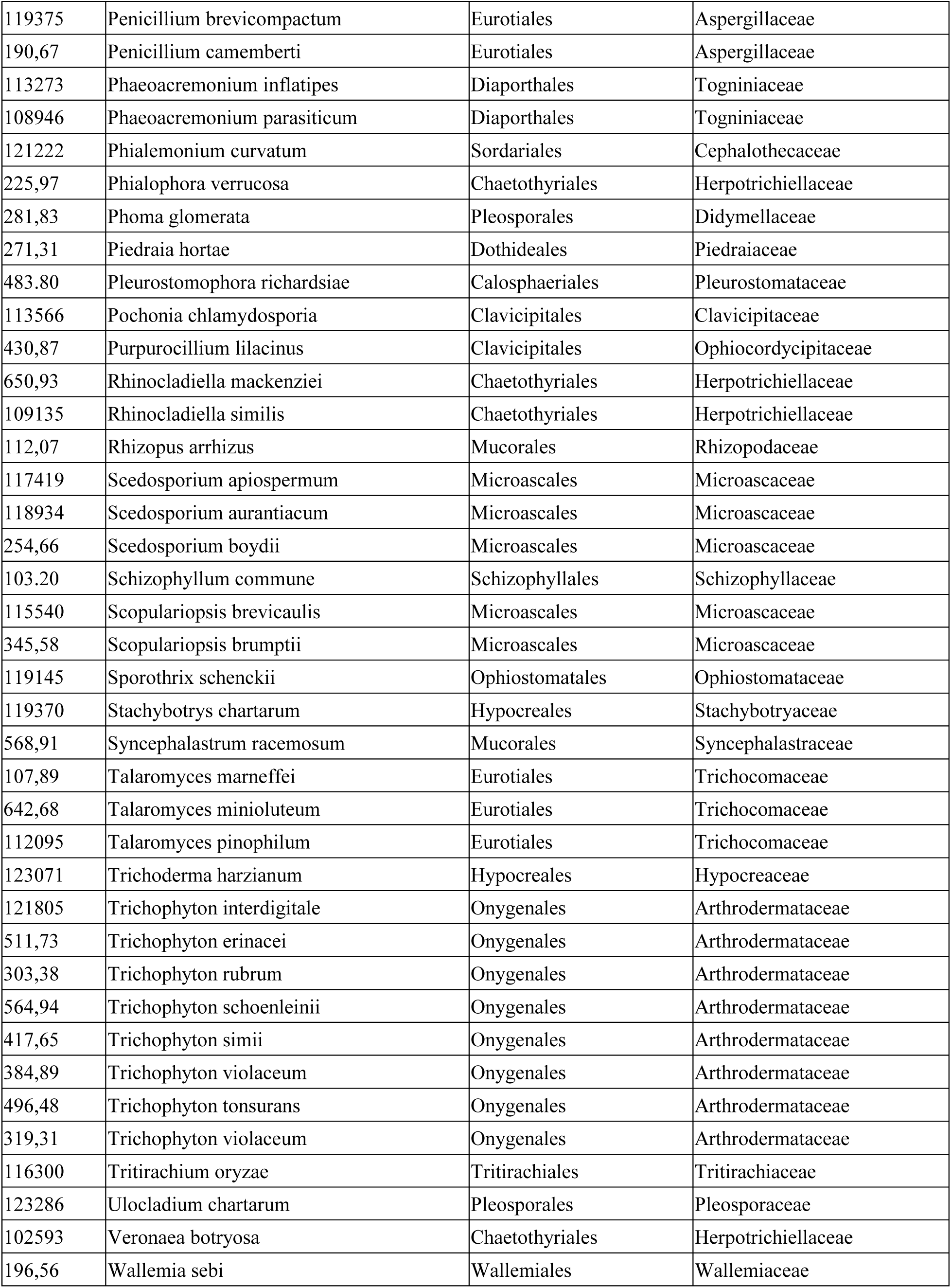
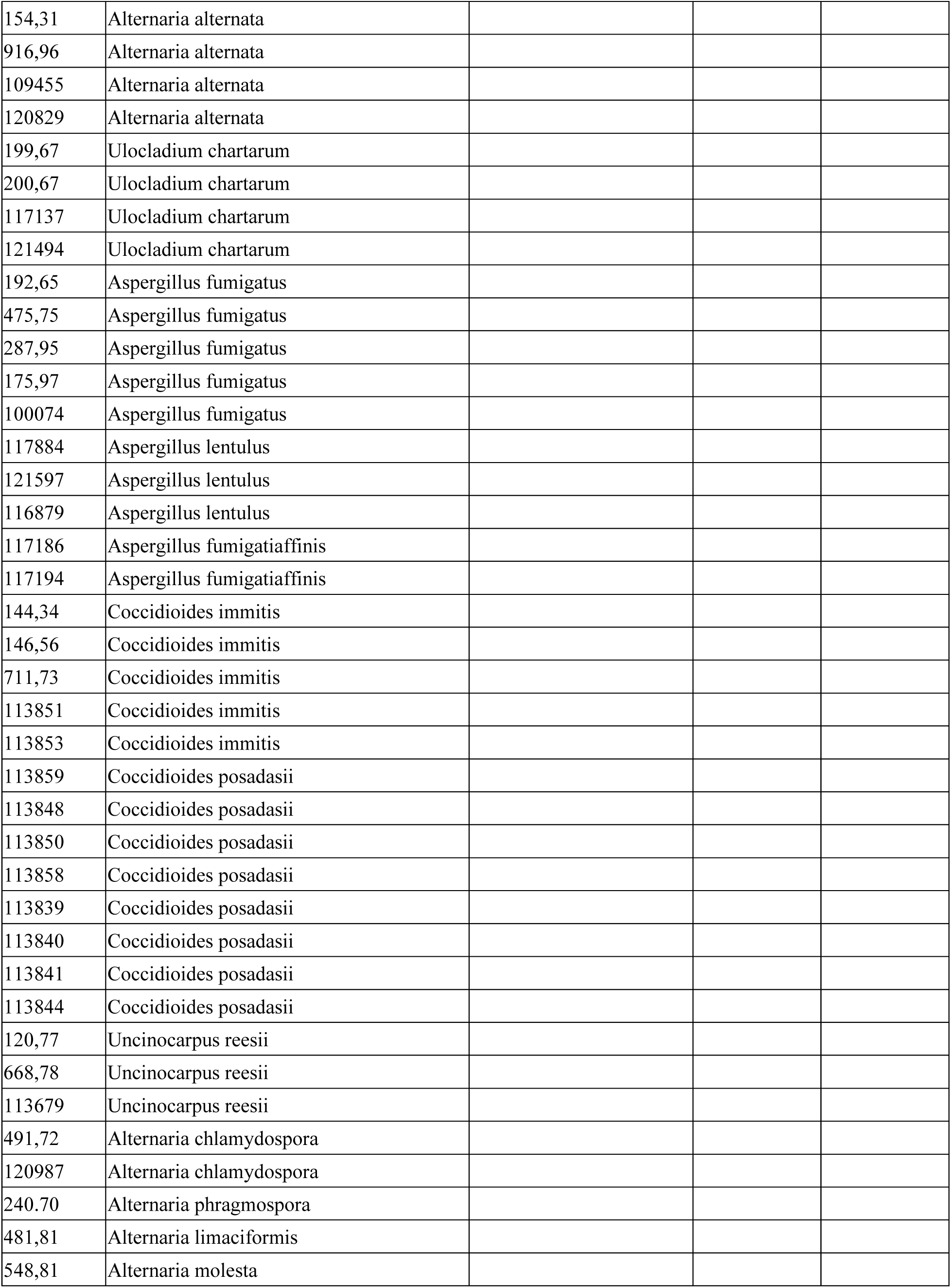
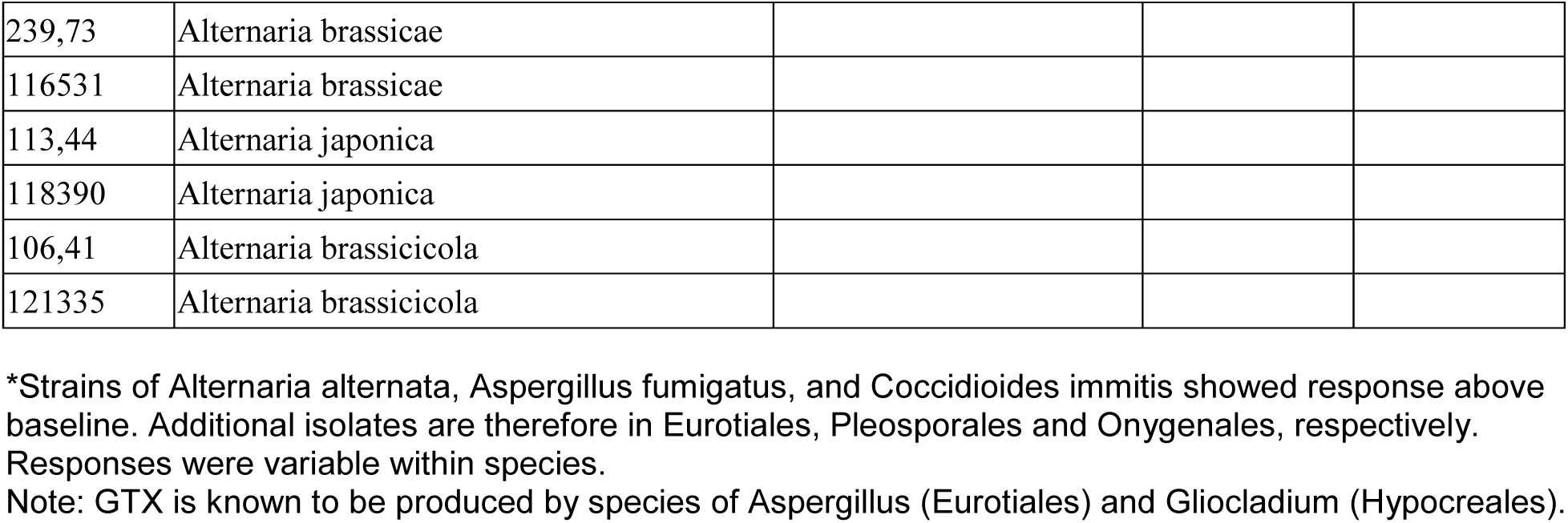
Strains analyzed for their ability to induce HIV-1 proviral expression.

| **CBS** | **Name** | **Order** | **Family** |
| --- | --- | --- | --- |
| 220,84 | *Acremonium recifei* | Hypocreales | Hypocreaceae |
| 603,78 | *Alternaria alternata* | Pleosporales | Pleosporaceae |
| 177,80 | *Alternaria caespitosa* | Pleosporales | Pleosporaceae |
| 210,86 | *Alternaria infectoria* | Pleosporales | Pleosporaceae |
| 163,74 | *Ascodesmis nigricans* | Pezizales | Ascodesmidaceae |
| 114,48 | *Aspergillus clavatus* | Eurotiales | Aspergillaceae |
| 625,66 | *Aspergillus flavus* | Eurotiales | Aspergillaceae |
| 113,26 | *Aspergillus fumigatus* | Eurotiales | Aspergillaceae |
| 416,68 | *Acremonium potronii* | Eurotiales | Aspergillaceae |
| 117884 | *Aspergillus lentulus* | Eurotiales | Aspergillaceae |
| 589,65 | *Aspergillus nidulans* | Eurotiales | Aspergillaceae |
| 769,97 | *Aspergillus niger* | Eurotiales | Aspergillaceae |
| 245,65 | *Aspergillus versicolor* | Eurotiales | Aspergillaceae |
| 116686 | *Aspergillus terreus* | Eurotiales | Aspergillaceae |
| 584,75 | *Aureobasidium pullulans* | Dothideales | Aureobasidiaceae |
| 172,57 | *Bipolaris australiensis* | Pleosporales | Pleosporaceae |
| 173,57 | *Bipolaris hawaiiensis* | Pleosporales | Pleosporaceae |
| 586,80 | *Bipolaris spicifera* | Pleosporales | Pleosporaceae |
| 113191 | *Botryosphaeria dothidea* | Botryosphaeriales | Botryosphaeriaceae |
| 567.78A | *Botrytis cinerea* | Helotiales | Sclerotiniaceae |
| 545,83 | *Chaetomium globosum* | Sordariales | Chaetomiaceae |
| 104,62 | *Chrysosporium keratinophilum* | Onygenales | Onygenaceae |
| 173,52 | *Cladophialophora bantiana* | Chaetothyriales | Herpotrichiellaceae |
| 316,56 | *Cladophialophora boppii* | Chaetothyriales | Herpotrichiellaceae |
| 166,54 | *Cladophialophora carrionii* | Chaetothyriales | Herpotrichiellaceae |
| 109501 | *Cladosporium cladosporioides* | Capnodiales | Cladosporiaceae |
| 399,80 | *Cladosporium bruhnei* | Capnodiales | Cladosporiaceae |
| 108,85 | *Cladosporium herbarum* | Capnodiales | Cladosporiaceae |
| 113856 | *Coccidioides immitis* | Onygenales | Onygenaceae |
| 113859 | *Coccidioides posadasii* | Onygenales | Onygenaceae |
| 117439 | *Coprinopsis cinereus* | Agaricales | Psathyrellaceae |
| 220,52 | *Curvularia geniculata* | Pleosporales | Pleosporaceae |
| 730,96 | *Curvularia lunata* | Pleosporales | Pleosporaceae |
| 637,82 | *Cylindrocarpon cyanescens* | Hypocreales | Hypocreaceae |
| 116632 | *Cyphellophora europaea* | Chaetothyriales | Cyphellophoraceae |
| 190,61 | *Cyphellophora laciniata* | Chaetothyriales | Cyphellophoraceae |
| 286,85 | *Cyphellophora pluriseptata* | Chaetothyriales | Cyphellophoraceae |
| 443,88 | Dissitimurus exedrus | Pleosporales | Pleosporaceae |
| 177,60 | Emmonsia crescens | Onygenales | Ajellomycetaceae |
| 102456 | Emergomyces pasteurianus | Onygenales | Ajellomycetaceae |
| 214,63 | Epidermophyton floccosum | Onygenales | Arthrodermataceae |
| 129,54 | Eurotium chevalieri | Eurotiales | Aspergillaceae |
| 207,35 | Exophiala dermatitidis | Chaetothyriales | Herpotrichiellaceae |
| 537,76 | Exophiala jeanselmei | Chaetothyriales | Herpotrichiellaceae |
| 899,68 | Exophiala spinifera | Chaetothyriales | Herpotrichiellaceae |
| 102455 | Exophiala xenobiotica | Chaetothyriales | Herpotrichiellaceae |
| 196,79 | Falciformispora senegalensis | Pleosporales | Leptosphaeriaceae |
| 102237 | Fonsecaea pedrosoi | Chaetothyriales | Herpotrichiellaceae |
| 840,88 | Fusarium oxysporum | Hypocreales | Hypocreaceae |
| 120996 | Fusarium proliferatum | Hypocreales | Hypocreaceae |
| 181,29 | Fusarium solani | Hypocreales | Hypocreaceae |
| 539,79 | Fusarium verticillioides | Hypocreales | Hypocreaceae |
| 109689 | Ganoderma lucidum | Polyporales | Polyporaceae |
| 224,48 | Geotrichum candidum | Saccharomycetales | Dipodascaceae |
| 136,72 | Histoplasma capsulatum | Onygenales | Ajellomycetaceae |
| 410,51 | Hortaea werneckii | Capnodiales | Capnodiaceae |
| 118183 | Hypoxylon investiens | Xylariales | Hypoxylaceae |
| 467,74 | Lomentospora prolificans | Microascales | Microascaceae |
| 331,50 | Madurella grisea | Sordariales | Chaetomiaceae |
| 110087 | Madurella mycetomatis | Sordariales | Chaetomiaceae |
| 119449 | Microsporum audouinii | Onygenales | Arthrodermataceae |
| 114329 | Microsporum canis | Onygenales | Arthrodermataceae |
| 427,63 | Microsporum ferrugineum | Onygenales | Arthrodermataceae |
| 246,89 | Monascus ruber | Eurotiales | Monascaceae |
| 157,58 | Moniliella suaveolens | Moniliellales | Moniliellaceae |
| 616,63 | Mucor racemosus | Mucorales | Mucoraceae |
| 120675 | Nannizzia gypsea | Onygenales | Arthrodermataceae |
| 116672 | Nannizziopsis vriesii | Onygenales | Nannizziopsidaceae |
| 204,33 | Neoscytalidium dimidiatum | Dothideales | Dothioraceae |
| 113,64 | Neosartorya fischeri | Eurotiales | Aspergillaceae |
| 331,78 | Neotestudina rosatii | Sordariales | Neotestudinaceae |
| 116660 | Ochroconis gallopava | Venturiales | Sympoventuriaceae |
| 109438 | Onychocola canadensis | Arachnomycetales | Arachnomycetaceae |
| 339,51 | Paecilomyces variotii | Eurotiales | Aspergillaceae |
| 272,86 | Penicillium funiculosum | Eurotiales | Aspergillaceae |
| 120336 | Penicillium chrysogenum | Eurotiales | Aspergillaceae |
| 119375 | *Penicillium brevicompactum* | Eurotiales | Aspergillaceae |
| 190,67 | *Penicillium camemberti* | Eurotiales | Aspergillaceae |
| 113273 | *Phaeoacremonium inflatipes* | Diaporthales | Togniniaceae |
| 108946 | *Phaeoacremonium parasiticum* | Diaporthales | Togniniaceae |
| 121222 | *Phialemonium curvatum* | Sordariales | Cephalothecaceae |
| 225,97 | *Phialophora verrucosa* | Chaetothyriales | Herpotrichiellaceae |
| 281,83 | *Phoma glomerata* | Pleosporales | Didymellaceae |
| 271,31 | *Piedraia hortae* | Dothideales | Piedraiaceae |
| 483.80 | *Pleurostomophora richardsiae* | Calosphaerales | Pleurostomataceae |
| 113566 | *Pochonia chlamydosporia* | Clavicipitales | Clavicipitaceae |
| 430,87 | *Purpureocillium lilacinus* | Clavicipitales | Ophiocordycipitaceae |
| 650,93 | *Rhinocladiella mackenziei* | Chaetothyriales | Herpotrichiellaceae |
| 109135 | *Rhinocladiella similis* | Chaetothyriales | Herpotrichiellaceae |
| 112,07 | *Rhizopus arrhizus* | Mucorales | Rhizopodaceae |
| 117419 | *Scedosporium apiospermum* | Microascales | Microascaceae |
| 118934 | *Scedosporium aurantiacum* | Microascales | Microascaceae |
| 254,66 | *Scedosporium boydii* | Microascales | Microascaceae |
| 103.20 | *Schizophyllum commune* | Schizophyllales | Schizophyllaceae |
| 115540 | *Scopulariopsis brevicaulis* | Microascales | Microascaceae |
| 345,58 | *Scopulariopsis brumptii* | Microascales | Microascaceae |
| 119145 | *Sporothrix schenckii* | Ophiostomatales | Ophiostomataceae |
| 119370 | *Stachybotrys chartarum* | Hypocreales | Stachybotryaceae |
| 568,91 | *Syncephalastrum racemosum* | Mucorales | Syncephalastraceae |
| 107,89 | *Talaromyces marneffeii* | Eurotiales | Trichocomaceae |
| 642,68 | *Talaromyces minioluteum* | Eurotiales | Trichocomaceae |
| 112095 | *Talaromyces pinophilum* | Eurotiales | Trichocomaceae |
| 123071 | *Trichoderma harzianum* | Hypocreales | Hypocreaceae |
| 121805 | *Trichophyton interdigitale* | Onygenales | Arthrodermataceae |
| 511,73 | *Trichophyton erinacei* | Onygenales | Arthrodermataceae |
| 303,38 | *Trichophyton rubrum* | Onygenales | Arthrodermataceae |
| 564,94 | *Trichophyton schoenleinii* | Onygenales | Arthrodermataceae |
| 417,65 | *Trichophyton simii* | Onygenales | Arthrodermataceae |
| 384,89 | *Trichophyton violaceum* | Onygenales | Arthrodermataceae |
| 496,48 | *Trichophyton tonsurans* | Onygenales | Arthrodermataceae |
| 319,31 | *Trichophyton violaceum* | Onygenales | Arthrodermataceae |
| 116300 | *Tritirachium oryzae* | Tritirachiales | Tritirachiaceae |
| 123286 | *Ulocladium chartarum* | Pleosporales | Pleosporaceae |
| 102593 | *Veronaea botryosa* | Chaetothyriales | Herpotrichiellaceae |
| 196,56 | *Wallemia sebi* | Wallemiales | Wallemiaceae |

|  |  |
| --- | --- |
| 154,31 | *Alternaria alternata* |
| 916,96 | *Alternaria alternata* |
| 109455 | *Alternaria alternata* |
| 120829 | *Alternaria alternata* |
| 199,67 | *Ulocladium chartarum* |
| 200,67 | *Ulocladium chartarum* |
| 117137 | *Ulocladium chartarum* |
| 121494 | *Ulocladium chartarum* |
| 192,65 | *Aspergillus fumigatus* |
| 475,75 | *Aspergillus fumigatus* |
| 287,95 | *Aspergillus fumigatus* |
| 175,97 | *Aspergillus fumigatus* |
| 100074 | *Aspergillus fumigatus* |
| 117884 | *Aspergillus lentulus* |
| 121597 | *Aspergillus lentulus* |
| 116879 | *Aspergillus lentulus* |
| 117186 | *Aspergillus fumigatiaffinis* |
| 117194 | *Aspergillus fumigatiaffinis* |
| 144,34 | *Coccidioides immitis* |
| 146,56 | *Coccidioides immitis* |
| 711,73 | *Coccidioides immitis* |
| 113851 | *Coccidioides immitis* |
| 113853 | *Coccidioides immitis* |
| 113859 | *Coccidioides posadasii* |
| 113848 | *Coccidioides posadasii* |
| 113850 | *Coccidioides posadasii* |
| 113858 | *Coccidioides posadasii* |
| 113839 | *Coccidioides posadasii* |
| 113840 | *Coccidioides posadasii* |
| 113841 | *Coccidioides posadasii* |
| 113844 | *Coccidioides posadasii* |
| 120,77 | *Uncinocarpus reesii* |
| 668,78 | *Uncinocarpus reesii* |
| 113679 | *Uncinocarpus reesii* |
| 491,72 | *Alternaria chlamydospora* |
| 120987 | *Alternaria chlamydospora* |
| 240,70 | *Alternaria phragmospora* |
| 481,81 | *Alternaria limaciformis* |
| 548,81 | *Alternaria molesta* |
| 239,73 | *Alternaria brassicae* |
| 116531 | *Alternaria brassicae* |
| 113,44 | *Alternaria japonica* |
| 118390 | *Alternaria japonica* |
| 106,41 | *Alternaria brassicicola* |
| 121335 | *Alternaria brassicicola* |

\*Strains of *Alternaria alternata*, *Aspergillus fumigatus*, and *Coccidioides immitis* showed response above baseline. Additional isolates are therefore in Eurotiales, Pleosporales and Onygenales, respectively. Responses were variable within species.
Note: GTX is known to be produced by species of *Aspergillus* (Eurotiales) and *Gliocladium* (Hypocreales).

**Supplementary Table 2.**
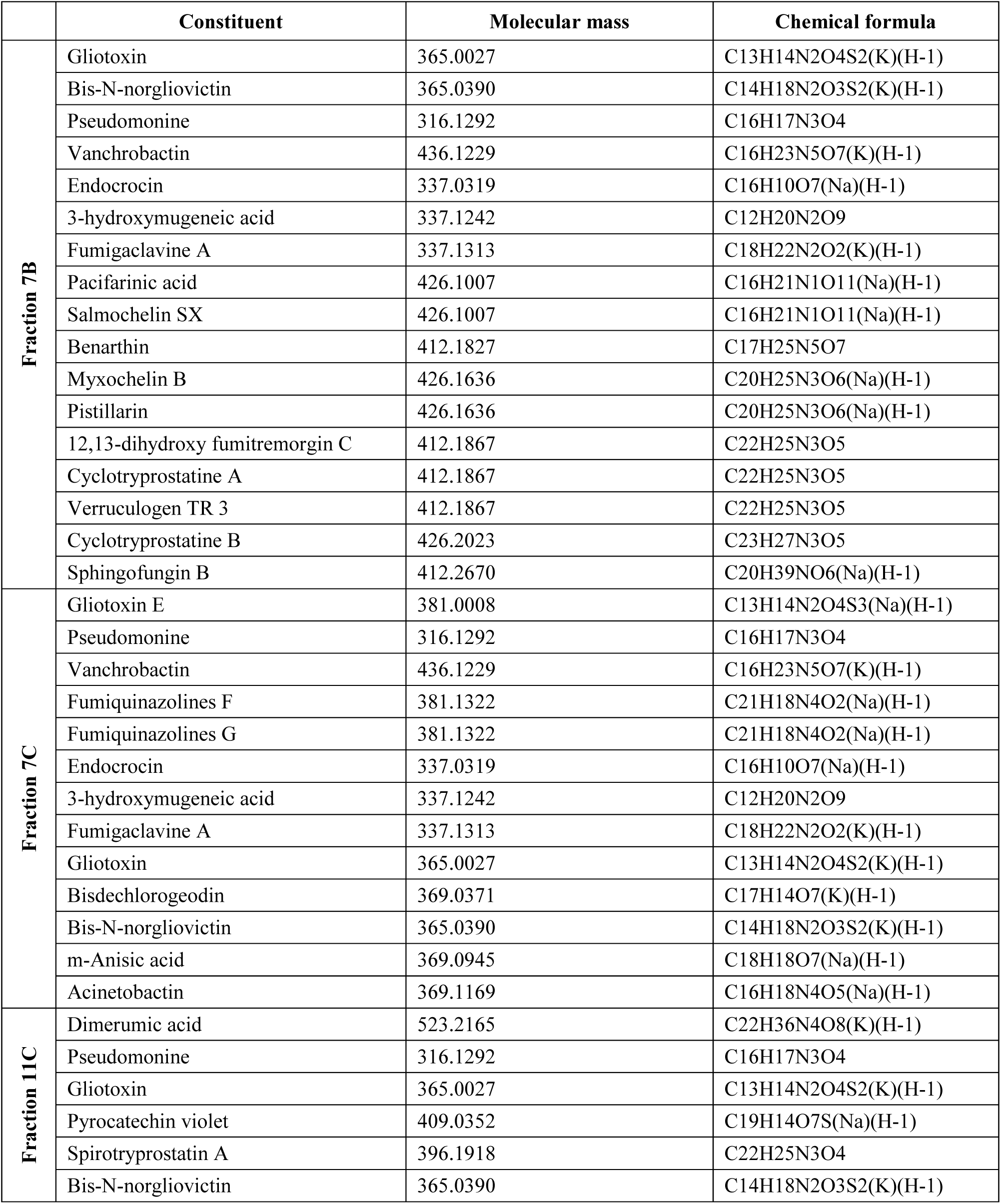
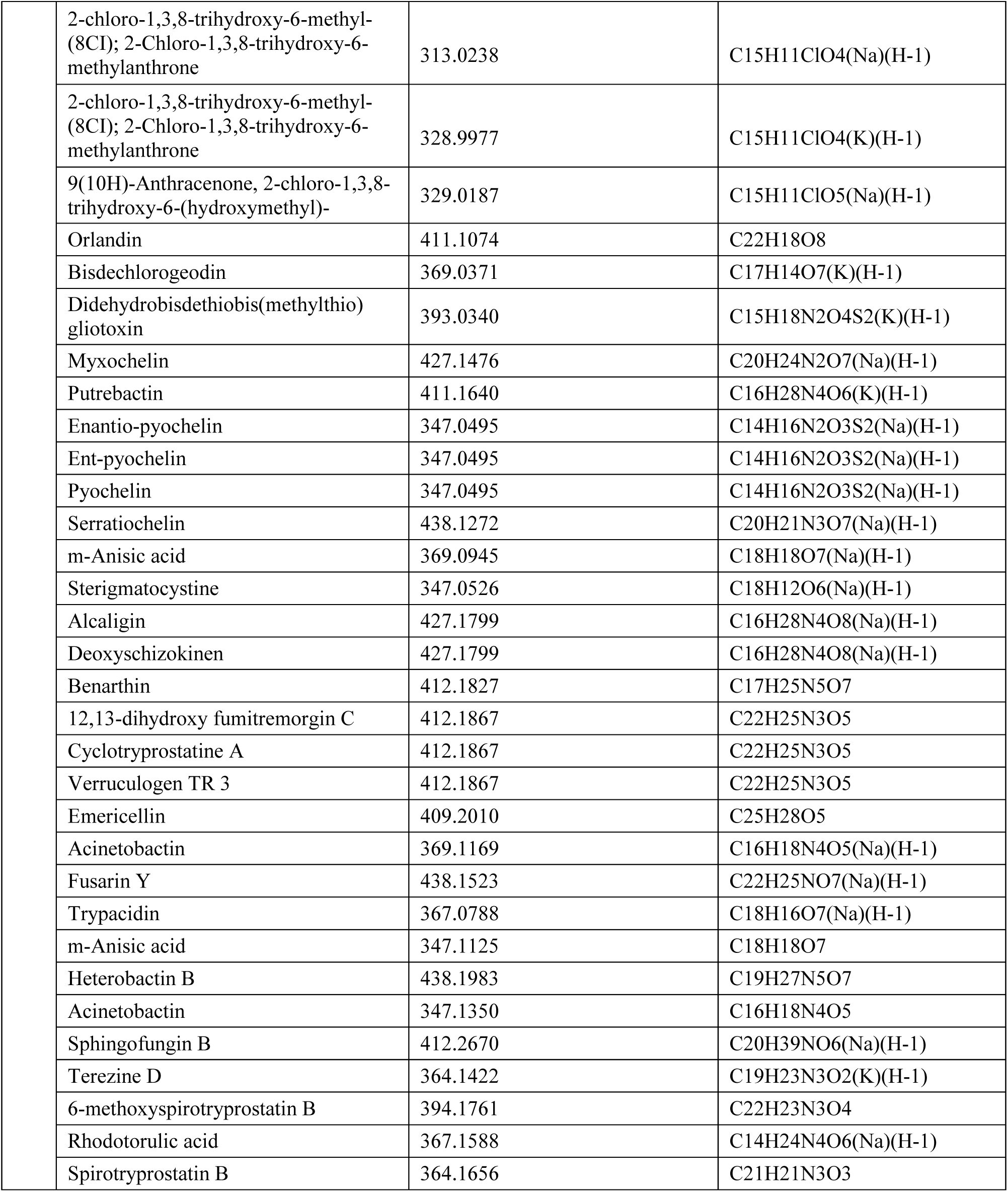
List of constituents of positive fractions annotated by MALDI-TOF mass spectrometry.

|  | Constituent | Molecular mass | Chemical formula |
| --- | --- | --- | --- |
| Fraction 7B | Gliotoxin | 365.0027 | C13H14N2O4S2(K)(H-1) |
|  | Bis-N-norgliovictin | 365.0390 | C14H18N2O3S2(K)(H-1) |
|  | Pseudomonine | 316.1292 | C16H17N3O4 |
|  | Vanchrobactin | 436.1229 | C16H23N5O7(K)(H-1) |
|  | Endocrocin | 337.0319 | C16H10O7(Na)(H-1) |
|  | 3-hydroxymugeneic acid | 337.1242 | C12H20N2O9 |
|  | Fumigaclavine A | 337.1313 | C18H22N2O2(K)(H-1) |
|  | Pacifarinic acid | 426.1007 | C16H21N1O11(Na)(H-1) |
|  | Salmoachelin SX | 426.1007 | C16H21N1O11(Na)(H-1) |
|  | Benarthin | 412.1827 | C17H25N5O7 |
|  | Myxochelin B | 426.1636 | C20H25N3O6(Na)(H-1) |
|  | Pistillarin | 426.1636 | C20H25N3O6(Na)(H-1) |
|  | 12,13-dihydroxy fumitremorgin C | 412.1867 | C22H25N3O5 |
|  | Cyclotryprostatine A | 412.1867 | C22H25N3O5 |
|  | Verruculogen TR 3 | 412.1867 | C22H25N3O5 |
|  | Cyclotryprostatine B | 426.2023 | C23H27N3O5 |
|  | Sphingofungin B | 412.2670 | C20H39NO6(Na)(H-1) |
| Fraction 7C | Gliotoxin E | 381.0008 | C13H14N2O4S3(Na)(H-1) |
|  | Pseudomonine | 316.1292 | C16H17N3O4 |
|  | Vanchrobactin | 436.1229 | C16H23N5O7(K)(H-1) |
|  | Fumiquinazolines F | 381.1322 | C21H18N4O2(Na)(H-1) |
|  | Fumiquinazolines G | 381.1322 | C21H18N4O2(Na)(H-1) |
|  | Endocrocin | 337.0319 | C16H10O7(Na)(H-1) |
|  | 3-hydroxymugeneic acid | 337.1242 | C12H20N2O9 |
|  | Fumigaclavine A | 337.1313 | C18H22N2O2(K)(H-1) |
|  | Gliotoxin | 365.0027 | C13H14N2O4S2(K)(H-1) |
|  | Bisdechlorogeodin | 369.0371 | C17H14O7(K)(H-1) |
|  | Bis-N-norgliovictin | 365.0390 | C14H18N2O3S2(K)(H-1) |
|  | m-Anisic acid | 369.0945 | C18H18O7(Na)(H-1) |
|  | Acinetobactin | 369.1169 | C16H18N4O5(Na)(H-1) |
| Fraction 11C | Dimerumic acid | 523.2165 | C22H36N4O8(K)(H-1) |
|  | Pseudomonine | 316.1292 | C16H17N3O4 |
|  | Gliotoxin | 365.0027 | C13H14N2O4S2(K)(H-1) |
|  | Pyrocatechin violet | 409.0352 | C19H14O7S(Na)(H-1) |
|  | Spirotryprostatin A | 396.1918 | C22H25N3O4 |
|  | Bis-N-norgliovictin | 365.0390 | C14H18N2O3S2(K)(H-1) |

|  |  |  |
| --- | --- | --- |
| 2-chloro-1,3,8-trihydroxy-6-methyl-(8Cl); 2-Chloro-1,3,8-trihydroxy-6-methylanthrone | 313.0238 | C15H11ClO4(Na)(H-1) |
| 2-chloro-1,3,8-trihydroxy-6-methyl-(8Cl); 2-Chloro-1,3,8-trihydroxy-6-methylanthrone | 328.9977 | C15H11ClO4(K)(H-1) |
| 9(10H)-Anthracenone, 2-chloro-1,3,8-trihydroxy-6-(hydroxymethyl)- | 329.0187 | C15H11ClO5(Na)(H-1) |
| Orlandin | 411.1074 | C22H18O8 |
| Bisdechlorogeodin | 369.0371 | C17H14O7(K)(H-1) |
| Didehydrobisdethiobis(methylthio)gliotoxin | 393.0340 | C15H18N2O4S2(K)(H-1) |
| Myxochelin | 427.1476 | C20H24N2O7(Na)(H-1) |
| Putrebactin | 411.1640 | C16H28N4O6(K)(H-1) |
| Enantio-pyochelin | 347.0495 | C14H16N2O3S2(Na)(H-1) |
| Ent-pyochelin | 347.0495 | C14H16N2O3S2(Na)(H-1) |
| Pyochelin | 347.0495 | C14H16N2O3S2(Na)(H-1) |
| Serratichelin | 438.1272 | C20H21N3O7(Na)(H-1) |
| m-Anisic acid | 369.0945 | C18H18O7(Na)(H-1) |
| Sterigmatocystine | 347.0526 | C18H12O6(Na)(H-1) |
| Alcaligin | 427.1799 | C16H28N4O8(Na)(H-1) |
| Deoxyschizokinen | 427.1799 | C16H28N4O8(Na)(H-1) |
| Benarthin | 412.1827 | C17H25N5O7 |
| 12,13-dihydroxy fumitremorgin C | 412.1867 | C22H25N3O5 |
| Cyclotryprostatine A | 412.1867 | C22H25N3O5 |
| Verruculogen TR 3 | 412.1867 | C22H25N3O5 |
| Emericellin | 409.2010 | C25H28O5 |
| Acinetobactin | 369.1169 | C16H18N4O5(Na)(H-1) |
| Fusarin Y | 438.1523 | C22H25NO7(Na)(H-1) |
| Trypacidin | 367.0788 | C18H16O7(Na)(H-1) |
| m-Anisic acid | 347.1125 | C18H18O7 |
| Heterobactin B | 438.1983 | C19H27N5O7 |
| Acinetobactin | 347.1350 | C16H18N4O5 |
| Sphingofungin B | 412.2670 | C20H39NO6(Na)(H-1) |
| Terezine D | 364.1422 | C19H23N3O2(K)(H-1) |
| 6-methoxyspirotryprostatin B | 394.1761 | C22H23N3O4 |
| Rhodotorulic acid | 367.1588 | C14H24N4O6(Na)(H-1) |
| Spirotryprostatin B | 364.1656 | C21H21N3O3 |

